# Working strokes produced by curling protofilaments at disassembling microtubule tips can be biochemically tuned and vary with species

**DOI:** 10.1101/2022.09.13.507824

**Authors:** Lucas E. Murray, Haein Kim, Luke M. Rice, Charles L. Asbury

**Affiliations:** Department of Physiology and Biophysics, University of Washington, Seattle, USA; Department of Biophysics, UT Southwestern Medical Center, Dallas, USA; Department of Biochemistry, UT Southwestern Medical Center, Dallas, USA; Department of Biochemistry, University of Washington, Seattle, USA

## Abstract

The disassembly of microtubules can generate force and drive intracellular motility. During mitosis, chromosomes remain persistently attached via kinetochores to the tips of disassembling microtubules, which pull the sister chromatids apart. According to the conformational wave hypothesis, such force generation requires that protofilaments curl outward from the disassembling tips to exert pulling force directly on kinetochores. Rigorously testing this idea will require modifying the mechanical and energetic properties of curling protofilaments, but no way to do so has yet been described. Here, by direct measurement of working strokes generated in vitro by curling protofilaments, we show that their mechanical energy output can be increased by adding magnesium, and that yeast microtubules generate larger and more energetic working strokes than bovine microtubules. Both the magnesium and species-dependent increases in work output can be explained by a lengthening of protofilament curls, without any change in their bending stiffness or intrinsic curvature. These observations demonstrate how work output from curling protofilaments can be tuned and suggest evolutionary conservation of the amount of curvature strain energy stored in the microtubule lattice.

## Introduction

Microtubules are filamentous polymers central to the active transport of cargoes in cells. While they often serve as passive tracks along which dynein and kinesin motors move, these filaments can also drive motility directly. Dynamic microtubules in the mitotic spindle transport chromosomes during cell division by shortening while their disassembling tips remain coupled via kinetochores to the chromosomes (Desai & Mitchison, 1997; Inoue & Salmon, 1995; McIntosh, Volkov, Ataullakhanov, & Grishchuk, 2010). Dynamic microtubules also generate force to properly position the mitotic spindle and the nucleus within cells (Carminati & Stearns, 1997; Dogterom, Kerssemakers, Romet-Lemonne, & Janson, 2005; Kozlowski, Srayko, & Nedelec, 2007; McIntosh et al., 2010; Nguyen-Ngoc, Afshar, & Gonczy, 2007). These microtubule-driven movements are powered by GTP hydrolysis. GTP is incorporated into the assembling polymer tip and then hydrolyzed, depositing energy into the GDP-tubulin lattice. The stored lattice energy is released during disassembly and can be harnessed to generate pulling force. Thus microtubules, like dynein and kinesin motors, convert chemical energy into mechanical work (McIntosh et al., 2010). How they do so remains poorly understood.

Two distinct classes of mechanism could explain how disassembling microtubule tips generate pulling force: the biased diffusion and conformational wave mechanisms. According to biased diffusion-based models, a tip-coupler such as the kinetochore undergoes a thermally driven random walk along the microtubule surface that is biased at the tip, due to the affinity of the coupler for the microtubule. If the affinity of the coupler for the microtubule is sufficiently high and if its diffusion is sufficiently fast, then the coupler can remain persistently associated with the disassembling tip, where it will experience a thermodynamic force in the direction of disassembly (Hill, 1985). The effect is analogous to capillary action that pulls fluids into narrow channels. Biased diffusion of a key kinetochore element, the Ndc80 complex, has been observed directly on microtubules in vitro (Powers et al., 2009).

By contrast, force generation in conformational wave-based models depends on structural changes at disassembling microtubule tips. During disassembly, individual rows of tubulin dimers called protofilaments curl outward from the tip before breaking apart, creating a wave of conformational change that propagates down the long axis of the microtubule (Kirschner, Williams, Weingarten, & Gerhart, 1974; Mandelkow & Mandelkow, 1985). These curling protofilaments are proposed to physically hook the kinetochore and pull against it to drive motility (Koshland, Mitchison, & Kirschner, 1988). Prior work showed that the amount of mechanical strain energy released by curling protofilaments is more than sufficient to account for kinetochore motility (Driver, Geyer, Bailey, Rice, & Asbury, 2017). However, whether kinetochores specifically harness any of this strain energy remains unclear, owing in part to the lack of methods for modifying mechanical or energetic properties of protofilament curls.

Many prior studies have established that added magnesium profoundly affects the dynamics of microtubules in vitro, altering the rates of switching between tip growth and shortening (Obrien, Salmon, Walker, & Erickson, 1990), accelerating tip disassembly (Martin, Butler, Clark, Zhou, & Bayley, 1987), and lengthening the protofilament curls at disassembling tips (Mandelkow, Mandelkow, & Milligan, 1991). Binding of magnesium to acidic residues in the disordered C-terminal tail of tubulin is implicated in magnesium-dependent acceleration of disassembly (Fees & Moore, 2018; Sackett, Bhattacharyya, & Wolff, 1985; Serrano, Avila, & Maccioni, 1984; Weisenberg, 1972). Faster disassembly by itself might explain why magnesium also lengthens protofilament curls, because it implies a faster rate of curling (i.e., that GDP-tubulins are losing their lateral bonds and curling outward more quickly) (Tran, Joshi, & Salmon, 1997). However, magnesium might also stabilize the longitudinal bonds within protofilament curls, thereby reducing the rate at which the curls break. To disentangle magnesium’s effects on curling and breakage rates, a systematic examination of curl contour length as a function of disassembly speed is required.

Previously we developed an assay for measuring forces and displacements generated by curling protofilaments (Driver et al., 2017) based on earlier pioneering work (Grishchuk, Molodtsov, Ataullakhanov, & McIntosh, 2005). In our “wave” assay, the curling protofilaments push laterally against a microbead tethered to the microtubule wall, thereby generating a brief pulse of bead motion against the force of a feedback-controlled laser trap. We show here that the sizes of these pulses – and the mechanical work energy that can be harnessed from them – are substantially increased by the addition of millimolar levels of magnesium. By measuring wave pulses after proteolytic cleavage of the β-tubulin C-terminal tail, we show that magnesium enlarges the pulses independently of its acceleration of disassembly, indicating that magnesium directly stabilizes the longitudinal bonds within protofilament curls. We also demonstrate that pulses generated by yeast tubulin are larger than those generated by bovine brain tubulin. A simple mechanical model shows that both the magnesium- and species-dependent changes in pulse energy can be explained solely by increasing the contour lengths of protofilament curls, without changing their intrinsic flexural rigidity or curvature. The conservation of protofilament flexural rigidity and stored lattice strain suggest that these biophysical properties are crucial to microtubule function in cells.

## Results

### Measuring outward curling of protofilaments from bovine brain microtubules

We previously measured the mechanical and energetic properties of protofilaments as they curled outward from recombinant yeast-tubulin microtubules (Driver et al., 2017). In our wave assay, a laser trap applies force against the curling protofilaments, via beads tethered to the microtubule lattice through a His_6_ tag on the C-terminus of β-tubulin (Johnson, Ayaz, Huddleston, & Rice, 2011). Linkage through the β-tubulin C-terminal tail creates a strong, flexible tether approximately 36 nm in length, which probably helps to avoid interference between the tethered bead and the curling protofilaments (Driver et al., 2017). To extend our approach to untagged mammalian brain tubulin, we modified the assay by introducing anti-His beads pre-decorated sparsely with the recombinant His_6_-tagged yeast tubulin into chambers containing coverslip-anchored microtubules growing from free bovine brain tubulin. The bead-linked yeast tubulin was incorporated into the assembling bovine microtubules, resulting in beads tethered to the sides of the filaments (Murray, Kim, Rice, & Asbury, 2022) (Figure 1a). Continuous tension, directed toward the plus end, was applied to a microtubule-tethered bead using feedback control. The tension pressed the bead against the microtubule lattice at a secondary contact point and suppressed Brownian motion, which facilitated tracking the bead with high spatiotemporal resolution. The microtubule plus end was then severed with laser scissors to induce disassembly (Franck, Powers, Gestaut, Davis, & Asbury, 2010). As the disassembling tip passed the secondary contact point, protofilament curls pushed laterally on the bead, causing it to rotate about its tether. This rotation produced a brief (100 – 400 ms) pulse of bead movement against the force of the laser trap, which was followed by bead detachment after further disassembly released the tether (Figure 1b). The pulses were parameterized by their amplitude relative to the baseline bead position (Figure 1–figure supplement 1), which is directly related to the lateral height that the protofilament curls project from the surface of the microtubule lattice (Figure 1–figure supplement 2) (Driver et al., 2017).

**Figure 1.**
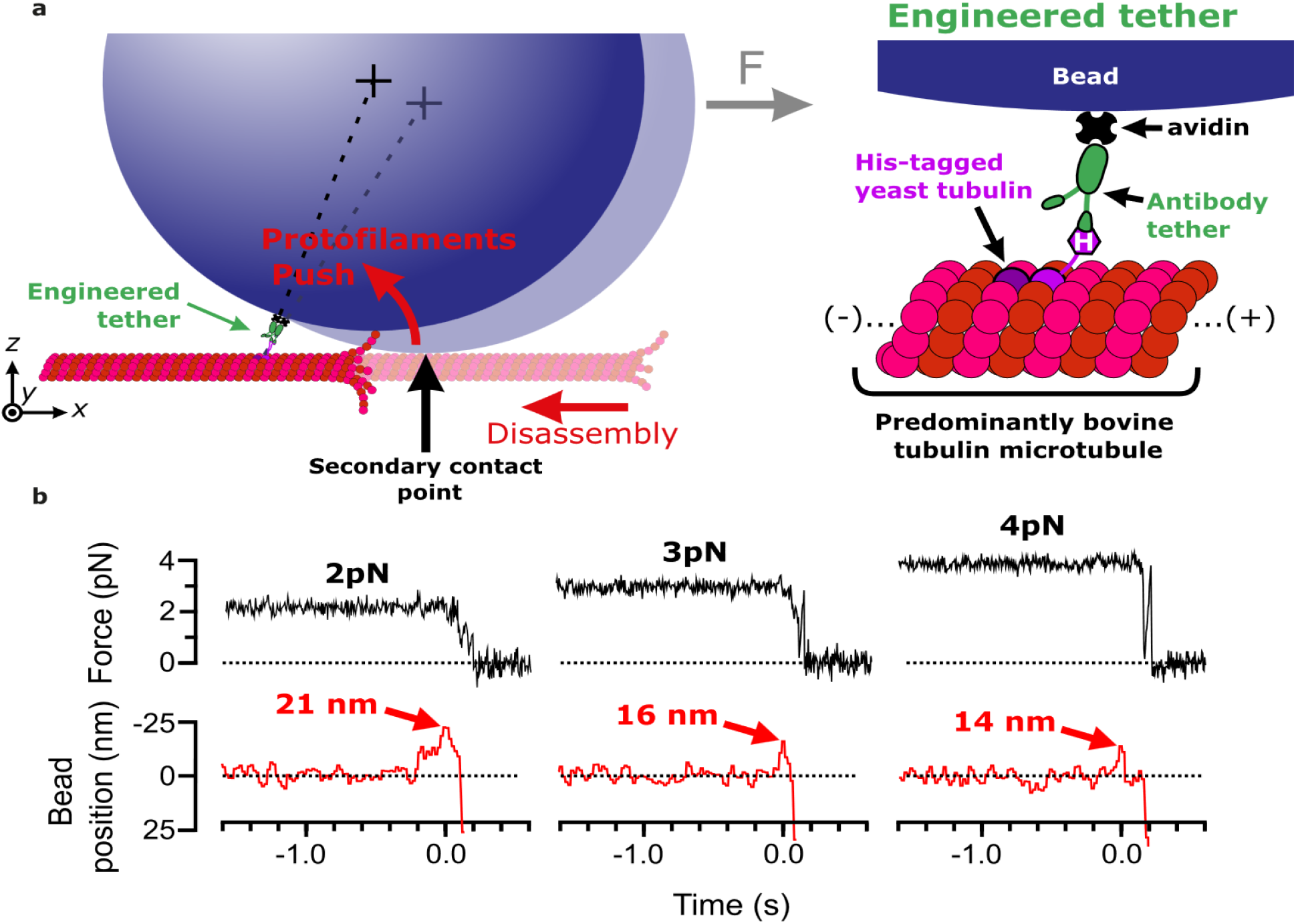
Measuring pulses of movement generated by protofilaments curling outward from the tips of disassembling bovine microtubules. (**a**) Schematic of the wave assay: A bead is tethered to the microtubule lattice via an engineered tether composed of recombinant His_6_-tagged yeast tubulin, a biotinylated anti-penta-His antibody, and streptavidin. Using a laser trap, the bead is tensioned towards the (+)-end, pressing it against the microtubule lattice at a secondary contact point. The stabilizing GTP cap is trimmed off the microtubule with laser scissors to initiate disassembly. Curling protofilaments at the disassembling microtubule tip form a conformational wave that pushes laterally on the bead, causing it to rock back about its tether. This rocking action produces a pulse of bead movement against the force of the laser trap. (**b**) Records of force (black) and bead position (red) versus time for three different bead-microtubule pairs. As the trapping force on the bead was increased, pulse heights decreased, consistent with spring-like behavior of the protofilament curls. *The following figure supplements are available for figure 1:* **Figure 1–figure supplement 1. Features of the pulses of bead motion generated by curling protofilaments.** **Figure 1–figure supplement 2. The bead and tether form a leverage system that amplifies protofilament curling motion.**

At 2 pN of trapping force, 59% of disassembly events yielded measurable pulses, with a mean amplitude of 18 nm. At higher forces, pulse amplitudes became smaller (Figure 1b), consistent with spring-like elasticity of the curling protofilaments, as we previously observed for yeast tubulin protofilament curls (Driver et al., 2017). Pulse amplitudes generated by bovine microtubules were smaller than those we measured previously from yeast microtubules at identical force levels (e.g., 19 versus 45 nm on average at 2 pN) (Driver et al., 2017). This observation suggests that bovine protofilament curls might be shorter than yeast curls, consistent with reports that disassembly products released from mammalian brain microtubules are shorter than their yeast-derived counterparts (Howes et al., 2018). Nevertheless, our findings confirm that pulses from bovine microtubules can be reliably measured using our modified wave assay.

### Adding magnesium enlarges the pulses generated by curling protofilaments

Divalent cations have long been known to affect tubulin self-association (Nogales et al., 1995; Olmsted & Borisy, 1975; Weisenberg, 1972), and to influence microtubule dynamics (Rosenfeld, Zackroff, & Weisenberg, 1976; Weisenberg, 1972). These effects are achieved partly through interactions of magnesium ions with the unstructured C-terminal tails of tubulin (Fees & Moore, 2018; Serrano, Delatorre, Maccioni, & Avila, 1984) and with the exchangeable and non-exchangeable nucleotide binding sites (Lee & Timasheff, 1975). Early cryo-electron microscopy of disassembling microtubules showed that magnesium lengthens protofilament curls at disassembling tips (Mandelkow et al., 1991). Based on these prior observations, we predicted that pulses recorded in our wave assay might become larger and more energetic with added magnesium.

As previously observed (Fees & Moore, 2018), we found that adding magnesium accelerated the disassembly of bovine brain tubulin microtubules, increasing their shortening speeds by about three-fold, from 430 nm·s^-1^ at our initial level of 1 mM magnesium to 1,200 nm·s^-1^ at 20 mM magnesium (Figure 2a and 2b). Consistent with our prediction, adding magnesium also increased the amplitudes of pulses measured in the wave assay (Figure 2c). At 2 pN of trapping force, the mean amplitude increased by 60% from 19 nm at 1 mM magnesium up to 29 nm at 20 mM magnesium (Figure 2d). This magnesium-dependent increase in pulse amplitude might be explained simply by a lengthening of the protofilament curls, as suggested by early cryo-electron microscopy studies. However, it might also reflect increases in the mechanical stiffness or curvature of the protofilaments, or in the number of protofilaments that push against the bead in the wave assay (as discussed below).

**Figure 2.**
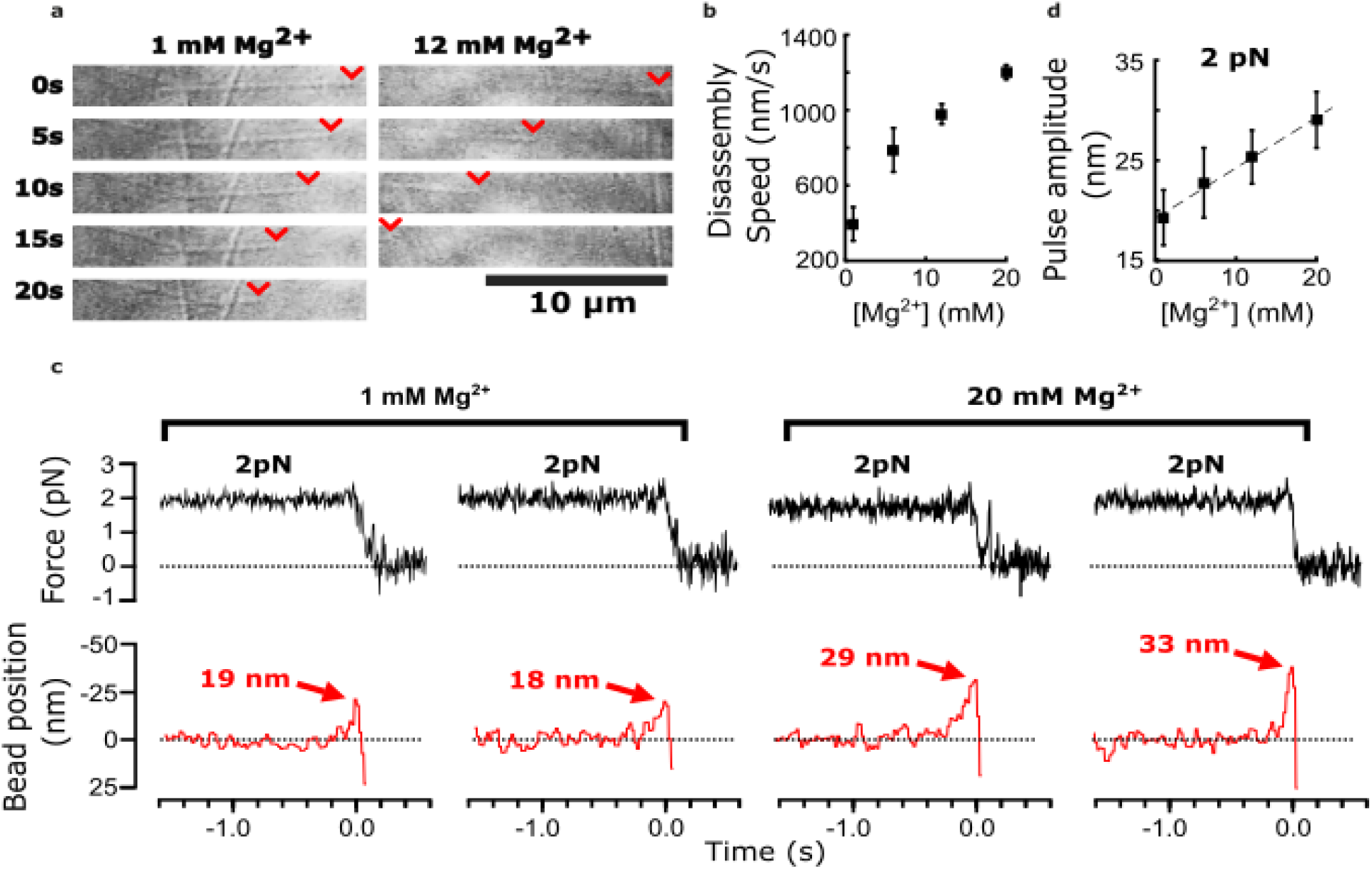
Added magnesium increases disassembly speed and pulse amplitude. (**a**) Time-lapse differential interference contrast images of individual microtubules disassembling in the presence of 1 or 12 mM magnesium. Arrowheads (red) indicate locations of disassembling tips. (**b**) Mean disassembly speed plotted against magnesium concentration. Error bars represent 95% confidence intervals, defined as ± (*t*·SEM), where *t* is drawn from Student’s *t*-distribution (with *ν* = *N* – 1 degrees of freedom and *N* = 34 – 51 samples per mean). (**c**) Records of force (black) and bead position (red) versus time for four bead-microtubule pairs, at two different magnesium concentrations. Pulse amplitudes were larger at the higher magnesium level. (**d**) Mean pulse amplitudes across four different magnesium concentrations, 1, 6, 12 and 20 mM. Error bars represent 95% confidence intervals (defined as in (b), with *ν* = *N* – 1 degrees of freedom and *N* = 25 – 40 samples per mean). Data in (c) and (d) were collected at 2 pN trap force.

### Adding magnesium increases work output from protofilament curls

To determine whether adding magnesium affects the mechanochemical work output from curling protofilaments, we measured pulse amplitudes across a variety of trapping forces and magnesium concentrations (Figure 3; and Figure 3–figure supplement 1). Measuring pulse amplitude as a function of force enables estimation of the total capacity for mechanical work output in the assay, which is given by the area under the amplitude-vs-force curve (Figure 3a) (Driver et al., 2017). Based on a line fit to the data, we estimated work output from the bovine brain microtubules in 1 mM magnesium at 105 pN·nm (Figure 3b). Adding magnesium increased the work output monotonically, raising it to 175 pN·nm at 20 mM magnesium (Figure 3b). This magnesium-induced increase was mainly due to enlargement of the pulses measured at low trapping force; extrapolating the line fits to zero force suggested that the unloaded pulse amplitude (i.e., the amplitude that would be measured in the absence of opposing trap force) increased 57% from 23 nm at 1 mM magnesium to 36 nm at 20 mM magnesium (Figure 3c). By contrast, extrapolating the linear fits to higher forces suggested relatively little change in maximum force at which the pulses were completely suppressed (~9 pN) (Figure 3a). Altogether, these observations show that magnesium increases mechanical work output from curling protofilaments while also increasing the lateral height that they project from the microtubule wall.

**Figure 3.**
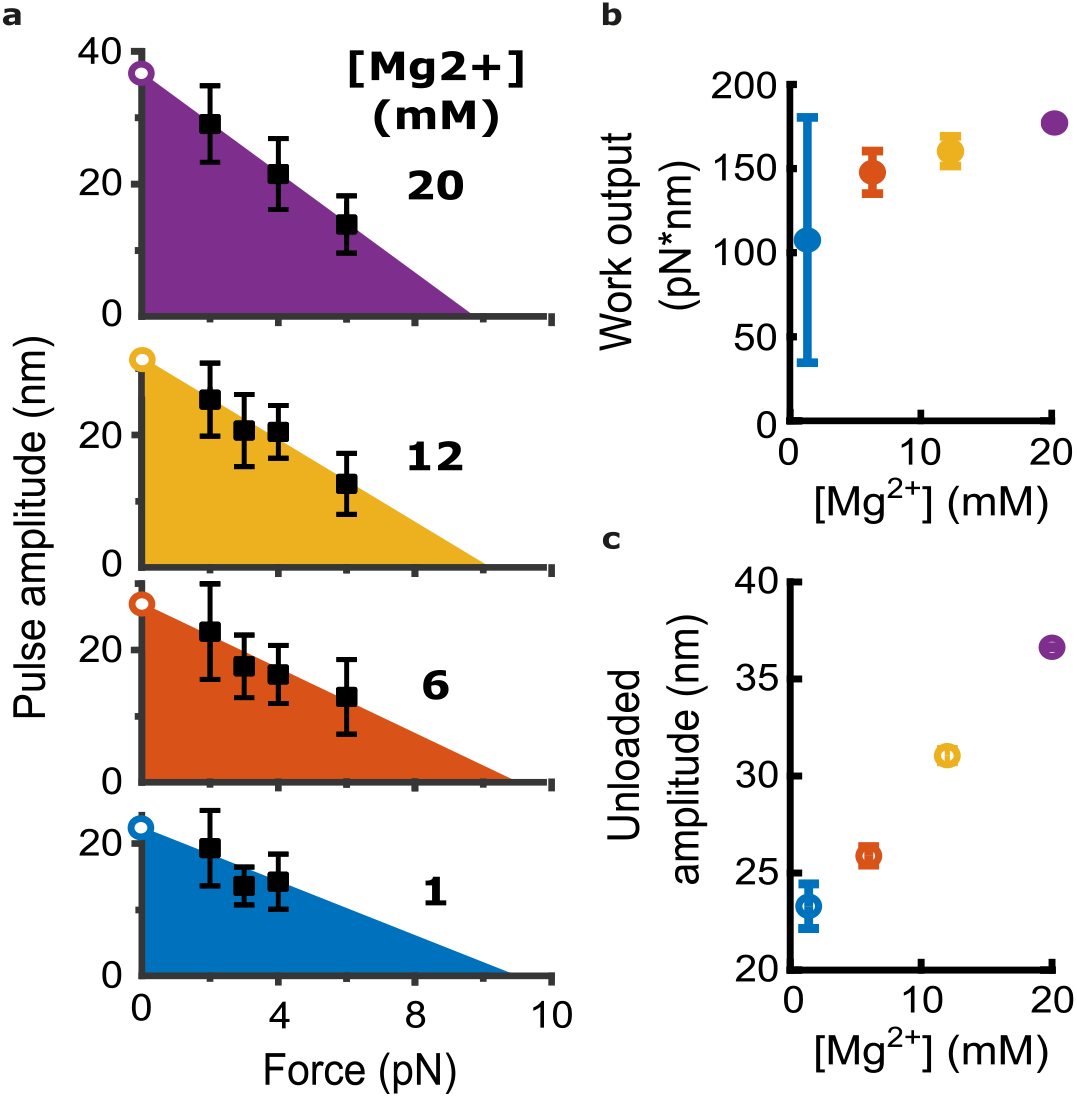
Magnesium increases the mechanical work output harnessed from curling protofilaments. (**a**) Mean pulse amplitudes (black squares) plotted against trapping force at the four indicated magnesium concentrations. Error bars represent 95% confidence intervals, defined as ± (*t*·SEM), where *t* is drawn from Student’s *t*-distribution (with *ν* = *N* – 1 degrees of freedom and *N* = 9 – 43 samples per mean). The capacity of protofilament curls to perform mechanical work in the assay was estimated at each magnesium concentration by fitting the amplitude versus force data with a line and then calculating the area under the line (colored triangular areas). To estimate unloaded pulse amplitudes, the line-fits were extrapolated to the y-intercept (open circles). (**b**) Mechanical work output, based on the colored areas shown in (a), plotted against magnesium concentration. Error bars represent 95% confidence intervals (estimated from the best-fit parameters, as explained in Materials & Methods). (**c**) Unloaded amplitudes, based on extrapolation of the line-fits in (a), plotted versus magnesium concentration. Error bars represent 95% confidence intervals (estimated as explained in Materials & Methods). *The following figure supplement is available for figure 3:* **Figure 3–figure supplement 1. Cumulative distributions of pulse amplitude measured with bovine tubulin at different trapping forces and magnesium levels.**

Notably, the mechanical work output from bovine microtubules was about three-fold less than we measured previously from microtubules composed entirely of recombinant yeast tubulin under similar conditions (~300 pN·nm at 1 mM magnesium) (Driver et al., 2017). This difference, like magnesium-dependent differences, could reflect altered contour lengths, bending stiffnesses, average curvatures, numbers of curling protofilaments pushing on the beads, or a combination thereof.

### Curl elongation alone can explain the magnesium-dependent increase in work output

To develop a deeper understanding of how magnesium increases the mechanical work output from curling protofilaments, we created a simple model of protofilament bending. The model relates structural aspects of protofilament curls, such as their relaxed curvature and the average number of dimers they contain, together with an estimate of their flexural rigidity, to predict the force-deflection behavior of a group of curls projecting radially outward from a microtubule tip. For simplicity, each individual tubulin dimer within a protofilament curl was modeled as a rigid rod connected to its longitudinal neighbors by Hookean bending springs (Figure 4a). These bending springs represented, in an idealized manner, all the contributions to elastic strain energy stored within a protofilament, including strain energy at the inter- and intra-dimer interfaces, and internal strain within the α- and β-tubulin core structures. Contour shapes for the individual protofilaments were solved by balancing the external force applied at their tip with the opposing bending spring torque at each inter-dimer node (Figure 4b, left). To model the force-deflection behavior of a group of protofilaments, single protofilaments were arranged radially, according to a 13-protofilament geometry (Figure 4b, right) (Amos & Klug, 1974). The bead was modeled as a rigid, flat surface since its curvature is negligible compared to that of the microtubule tip. Prior cryo-electron tomography studies of disassembling microtubules found almost all the variation in protofilament shape to occur in the radial direction (i.e., within a plane coincident with both the relaxed contour and the long axis of the microtubule) (McIntosh et al., 2018). Therefore, protofilament bending in our model was limited to the radial direction.

**Figure 4.**
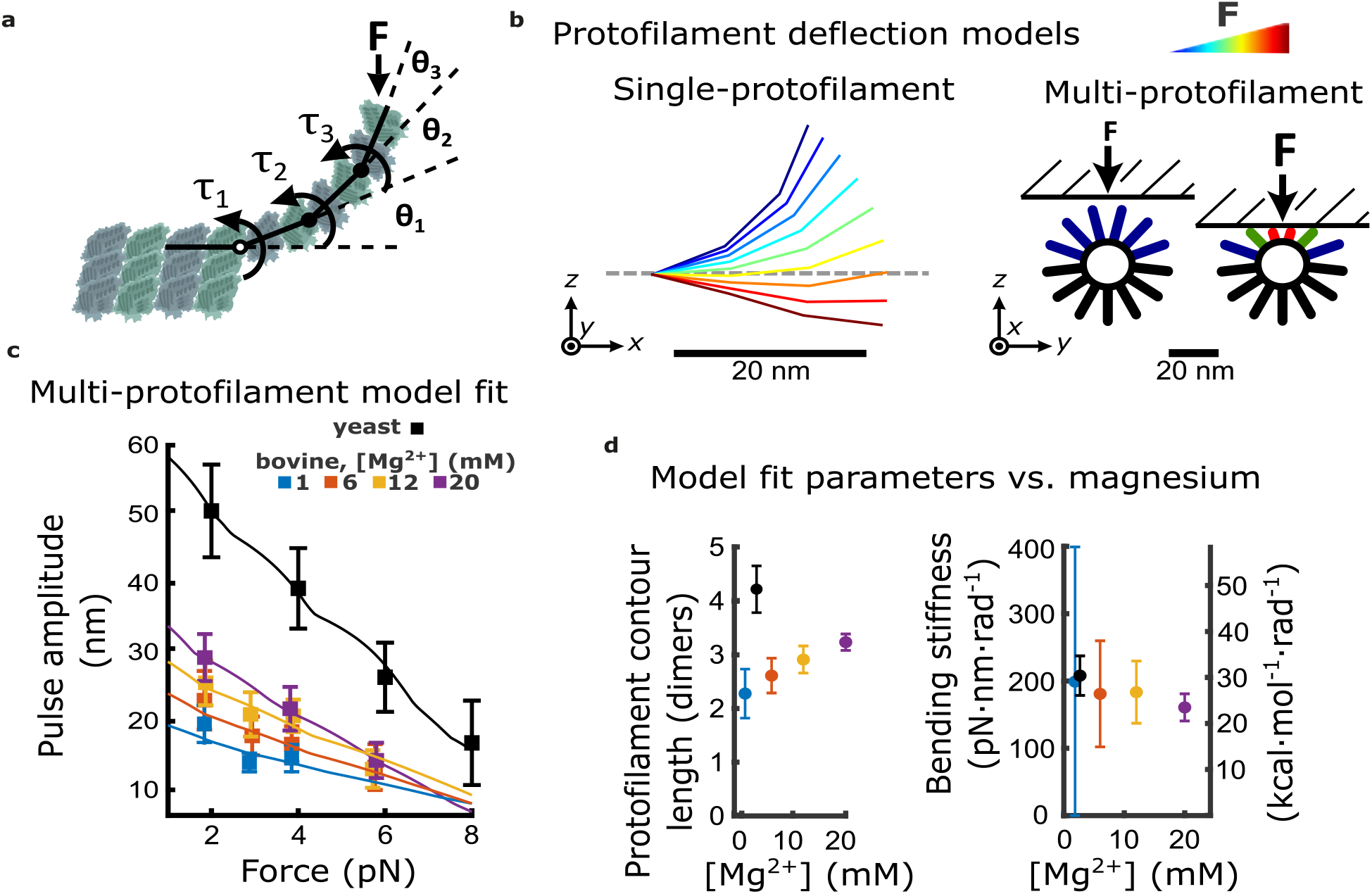
Magnesium- and species-dependent increases in work output can be explained solely by a lengthening of protofilament curls. (**a**) Model for bending of a single protofilament. Tubulin dimers are represented as rigid rods linked by Hookean torsion springs with relaxed angles of 23°. The springs are simplified representations of all contributions to elastic strain energy storage within a protofilament, including at intra- and inter-dimer interfaces, and within the core α- and β-tubulin structures. An external force, *F*, perpendicular to the microtubule long-axis, is applied at the protofilament tip. The balance between *F* and the torsion at each bending node, *τ_n_*, is used to calculate the contour shape of the protofilament (i.e., the angles *θ_n_*). (**b**) Calculated shapes for a single protofilament at different levels of external force (indicated by the color legend). Model for deflection of multiple protofilaments at a microtubule tip, seen end-on. Single protofilaments, modeled as in (a), are arranged radially according to the geometry of a 13-protofilament microtubule. The bead is modeled as a flat rigid surface, pushed downward onto the protofilaments to predict a force-deflection relationship. Cartoon at right shows distribution of protofilament deflections for an arbitrary bead height. (**c**) Amplitude versus force curves predicted by the multi-protofilament model, after fitting to measured pulse data (symbols) at indicated magnesium concentrations. Bovine data are recopied from Figure 3a. Yeast data combine new measurements with data previously published in (Driver, 2017). (**d**) Two fitted parameters, the mean contour length and bending stiffness (flexural rigidity) of protofilament curls, plotted versus magnesium concentration. The fitted contour length increases with added magnesium, and is larger for yeast microtubules, while the apparent flexural rigidity remains unchanged. *The following figure supplements are available for figure 4:* **Figure 4—figure supplement 1: How force-deflection behavior of the single protofilament model changes with variation in the number of segments (dimers), the intrinsic bending stiffness, and the relaxed angle per tubulin dimer.** **Figure 4—figure supplement 2: Table of estimates of protofilament curvature reported in the literature.** **Figure 4—figure supplement 3: Estimates of protofilament curvature from micrographs of disassembling tips presented in Mandelkow et al. 1991.** **Figure 4—figure supplement 4: Comparison of force-deflection relationship for a single protofilament, and multiple protofilaments arranged to reflect geometry at a microtubule tip.** **Figure 4—figure supplement 5: Comparison of disassembly speeds for bovine versus yeast microtubules.**

Given these assumptions, deflection of individual protofilaments varied according to their orientation relative to the bead surface (Figure 4b, right). A detailed analysis of changes in the force-deflection profile that occur with respect to changes in the average curvature, average dimers per curl, and flexural rigidity is shown in the supplemental material (Figure 4–figure supplement 1).

To fit the behavior of this multi-protofilament model to the measured pulse amplitude versus force data at each magnesium concentration, we adjusted the average number of dimers in each curl (i.e., the curl contour length) and the stiffness of the bending springs. We kept the relaxed angle per dimer fixed at 23° because, in the absence of MAPs, the curvature of protofilaments at microtubule tips disassembling in vitro is consistently between 20° and 25° per dimer (Figure 4–figure supplement 2) and this curvature does not change appreciably with added magnesium (Figure 4–figure supplement 3) (Mandelkow et al., 1991) or calcium (Muller-Reichert, Chretien, Severin, & Hyman, 1998). Because the bead acts as a lever, measured axial displacements of the bead are larger than the lateral deflections of the protofilaments by a leverage factor of approximately 2-fold (Figure 1–figure supplement 2) (Driver et al., 2017). Predicted amplitude-vs-force curves were roughly linear, but with slight “ripples” that occurred because movement of the bead toward the microtubule gradually engaged more protofilaments (Figure 4c; Figure 4–figure supplement 4) (See Materials & Methods for details). Optimal fit parameters are plotted as functions of magnesium in Figure 4d.

The fitted contour lengths of protofilaments increased monotonically with added magnesium, from 2.3 ± 0.5 dimers at 1 mM magnesium to 3.2 ± 0.2 dimers at 20 mM. However, the fitted bending stiffness per dimer, 175 pN·nm·rad^-1^, did not appreciably change with added magnesium (Figure 4d). These results suggest that magnesium increases pulse amplitude and work output by lengthening protofilament curls, without eliciting any change in their intrinsic stiffness or curvature.

### Curl elongation alone explains the larger pulses from yeast microtubules

To understand why yeast microtubules generated larger, more energetic pulses relative to bovine microtubules, we fit our multi-protofilament model to the amplitude versus force data measured from microtubules composed entirely of recombinant yeast tubulin (Figure 4c). As in our analysis of the bovine microtubule data, we allowed both the curl contour length and the stiffness of the bending springs to vary while keeping the relaxed angle per dimer fixed at 23°, consistent with cryo-ET tomograms of kinetochore microtubules in yeast (McIntosh et al., 2018). The contour length that best fit the yeast data, 4.4 ± 0.5 dimers per curl, was 1.9-fold higher than the contour length inferred at identical magnesium concentration (1 mM) from the bovine data, 2.3 ± 0.5 dimers per curl (Figure 4d). The bending stiffness per dimer that best fit the yeast data, 206 ± 44 pN·nm·rad^-1^, was statistically indistinguishable from that inferred from the bovine data (Figure 4d). These observations suggest that protofilament curls at yeast microtubule tips are longer but have the same intrinsic mechanical rigidity as the curls at bovine microtubule tips.

### Removing the β-tubulin tail suppresses magnesium’s enhancement of disassembly speed but not of pulse amplitude

Prior studies have suggested that longer protofilament curls might arise simply as a consequence of faster disassembly speeds (Tran et al., 1997). Consistent with this view, when we increased magnesium from 1 to 20 mM we observed a 3-fold increase in disassembly speed (Figure 2b) concomitant with a 1.6-fold increase in pulse amplitude (Figures 2d and 4c). Likewise, yeast microtubules disassembled 4-fold faster than bovine microtubules at 1 mM magnesium (Figure 4—figure supplement 5) and generated 3-fold larger pulses (Figure 4c). Faster disassembly speeds imply that GDP-tubulins lose their lateral bonds more quickly, which equivalently can be viewed as an accelerated rate of growth of the protofilament curls at disassembling tips. However, curl size is dictated not only by curl growth but also by curl breakage; steady state curl length will depend on a kinetic balance between the rates of curling and breakage (Tran et al., 1997). In principle, both these rates could vary in a magnesium-dependent manner. To distinguish the potential influence of magnesium on curl breakage from its obvious effect on disassembly speed (and therefore on curl growth rate), we sought a method to slow bovine microtubule disassembly at elevated levels of magnesium. A recent discovery pointed to one such method. Fees & Moore found that removing the β-tubulin C-terminal tail, by treating microtubules with the protease subtilisin, suppresses the effect of magnesium on disassembly speed (Fees & Moore, 2018). Thus, at high magnesium concentration, subtilisin-treated microtubules disassemble much more slowly than untreated microtubules. If magnesium lengthens protofilament curls solely because it accelerates disassembly, then subtilisin treatment should suppress the magnesium-dependent enlargement of pulses in the wave assay.

Contrary to this prediction, however, subtilisin treatment did not reduce pulse amplitudes in the wave assay. Consistent with the prior work of Fees and Moore, we found that subtilisin treatment for 10 to 20 minutes was sufficient to remove the β-tubulin C-terminal tail (Figure 5a) and to suppress almost completely the magnesium-dependent acceleration of disassembly (Figure 5b). While disassembly of untreated control microtubules was strongly accelerated, from 400 to 1,200 nm·s^-1^, when magnesium was increased from 1 to 20 mM, the disassembly after 10 or more minutes of subtilisin treatment was consistently slower and remained at approximately 300 nm·s^-1^ across the same range of magnesium levels. Despite this strikingly slower disassembly, the mean pulse amplitude measured in the wave assay after subtilisin-treatment remained at least as large as that measured in controls with untreated tubulin (Figure 5c). At 20 mM magnesium, the mean pulse amplitude generated after 5 to 10 min of subtilisin treatment was 30 to 40 nm (respectively), a size very similar to (or even slightly larger than) the mean amplitude generated by untreated microtubules, which was 28 nm. This observation indicates that magnesium enlarges protofilament curls independently of its acceleration of disassembly and suggests that distinct magnesium-binding sites probably underlie these two effects.

**Figure 5.**
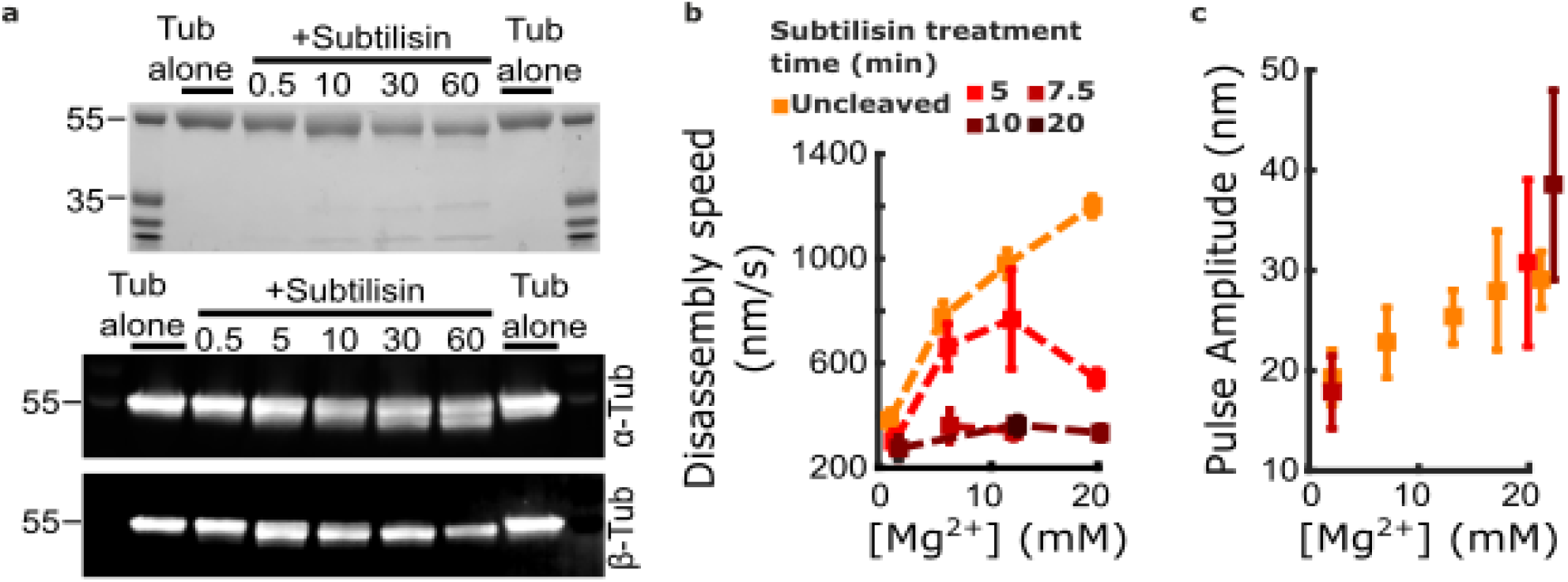
Removing the β-tubulin C-terminal tail suppresses magnesium’s acceleration of disassembly speed but not its enhancement of pulse amplitude. (**a**) Proteolytic products of tubulin treated with subtilisin for the indicate times (in minutes), before quenching with 1 mM PMSF, visualized by Coomassie staining (top image) or Western blotting (bottom image). αTub DM1A:FITC mouse conjugate 1:1000 and βTub CST 9F3 rabbit 1:1000 were used for primary antibody staining. (**b**) Mean disassembly speeds, measured after treatment of tubulin with subtilisin for the indicated durations and plotted versus magnesium concentration. The large magnesium-dependent acceleration of disassembly seen with untreated tubulin (orange symbols) was suppressed after 10 – 20 min subtilisin treatment (dark red symbols). Error bars represent 95% confidence intervals, defined as ± (*t*·SEM), where *t* is drawn from Student’s *t*-distribution (with *ν* = *N* – 1 degrees of freedom and *N* = 5 – 51 samples per mean). Data for untreated tubulin recopied from Figure 2b. (**c**) Pulse amplitudes, measured in the wave assay at 2 pN trapping force after treatment of tubulin with subtilisin, plotted versus magnesium concentration. Symbol colors indicate subtilisin treatment times according to the legend of (a). Treatment with subtilisin did not suppress the effect of magnesium on pulse amplitude. Error bars represent 95% confidence intervals, defined as ± (*t*·SEM), where *t* is drawn from Student’s *t*-distribution (with *ν* = *N* – 1 degrees of freedom and *N* = 28 – 44 samples per mean).

## Discussion

### Yeast and mammalian microtubules store similar lattice strain energies

Fitting wave assay data with the multi-protofilament model has allowed us to directly estimate a key biophysical property of curling protofilaments at disassembling microtubule tips: their flexural rigidity. Previously, this property was only inferred indirectly, from static cryo-electron tomograms (McIntosh et al., 2018), or from stiffness measurements of intact microtubules (Hawkins, Mirigian, Yasar, & Ross, 2010; Kononova et al., 2014) (Molodtsov, Grishchuk, Efremov, McIntosh, & Ataullakhanov, 2005; VanBuren, Cassimeris, & Odde, 2005). Our fitted estimate for bending stiffness, 175 pN·nm·rad^-1^, implies that fully straightening a protofilament from its relaxed curvature into a lattice-compatible state would require approximately 17 k_B_T, or 10 kcal/mol of work energy per tubulin dimer. This represents a very substantial fraction (^~^80%) of the free energy available from GTP hydrolysis, ^~^12.3 kcal/mol (Desai & Mitchison, 1997; Howard, 1996), consistent with previous suggestions that most of the energy derived from hydrolysis is stored as curvature strain in the microtubule lattice (Caplow, Ruhlen, & Shanks, 1994), and consistent with our previous lower-bound estimate (Driver et al., 2017). Moreover, our analysis suggests that the flexural rigidity of curling protofilaments is conserved between yeast and bovine tubulin and therefore that the amount of strain energy stored per tubulin dimer in the microtubule lattice is probably also conserved.

The idea that protofilament flexural rigidity and stored lattice strain are conserved, despite a billion years of evolution separating yeast and vertebrates, suggests that these biophysical properties are crucial to microtubule function. Indeed, most current models assume that microtubule dynamic instability arises from the counteracting influences of lateral bonding versus lattice strain, which tend to stabilize and destabilize the polymer, respectively (Gudimchuk et al., 2020; McIntosh et al., 2018; VanBuren et al., 2005; VanBuren, Odde, & Cassimeris, 2002). Given the importance of dynamic instability for cell viability, there may be strong selective pressure to maintain a specific lattice strain energy.

### Protofilament curl length can affect mechanical work output

In contrast to their consistent flexural rigidity, the average length of protofilament curls at disassembling microtubule tips can vary widely depending on tubulin species and buffer conditions (McIntosh et al., 2018; McIntosh et al., 2013). By our estimates, the average curl length grew ^~^50% as magnesium was increased from 1 to 20 mM. And curls at yeast microtubule tips were ^~^2-fold larger than those at bovine microtubule tips. These curl enlargements were associated with greater mechanical work output in the wave assay, as expected, since longer curls store more elastic energy and can push the bead laterally farther away from the microtubule surface. Whether longer protofilament curls can similarly enhance microtubule-driven motility in vivo remains unknown. In budding yeast, where each kinetochore attaches a single microtubule tip (Winey et al., 1995) and robust tip-coupling depends on the ring-forming Dam1 complex (Miranda, De Wulf, Sorger, & Harrison, 2005; Umbreit et al., 2014; Westermann et al., 2005; Westermann et al., 2006), a minimum curl length might be required for a microtubule-encircling Dam1 ring to efficiently harness curl energy via the conformational wave mechanism (Molodtsov et al., 2005). However, beyond a certain length, further enlargement of the curls might not be beneficial. In other species whose kinetochores attach numerous microtubule tips and lack any ring-forming complexes, coupling might depend less on the conformational wave mechanism and instead might rely on biased diffusion. An attractive idea is that the larger and more energetic pulses produced by yeast microtubules in the wave assay, as compared to bovine microtubules, might reflect stronger selective pressure to maintain long protofilament curls in yeast, because yeast might depend more heavily on long curls for mitosis. Alternatively, plus end-binders (+TIPs) and other microtubule-associated proteins that alter microtubule tip morphology (Cassimeris, Gard, Tran, & Erickson, 2001; Desai, Verma, Mitchison, & Walczak, 1999; Farmer, Arpa, Hall, & Zanic, 2021; Girao et al., 2020; Kerssemakers et al., 2006) could enlarge (or stiffen) protofilament curls, potentially enhancing their mechanical work output in a spatiotemporally regulated manner.

### Magnesium directly inhibits the breakage of protofilament curls

Our measurements also reveal new information about the relationship between disassembly speed and protofilament curl length, and about the mechanisms by which magnesium affects these tip properties. Classic work suggested that elongation of protofilament curls by magnesium might be a simple indirect consequence of its acceleration of disassembly (Tran et al., 1997). However, in 20 mM magnesium, subtilisin-treated microtubules disassembled three-fold more slowly than untreated controls and yet their pulse amplitudes remained consistently elevated, at well over 30 nm on average. Likewise, we previously showed that microtubules composed of a hyperstable T238V mutant tubulin disassemble seven-fold more slowly and yet they generated pulses with amplitudes indistinguishable from wild-type (Driver et al., 2017). These observations indicate that disassembly speed and curl length are not strictly coupled, and that magnesium-dependent enlargement of protofilament curls is not simply a consequence of accelerated disassembly. Rather, magnesium must directly inhibit the breakage of protofilament curls.

The effects of magnesium on disassembly speed and on curl length appear to be mediated by different interaction sites on tubulin. Magnesium’s acceleration of disassembly depends on the β-tubulin C-terminal tail, since this effect is suppressed upon removal of the β-tail by subtilisin (Fees & Moore, 2018). But subtilisin treatment did not suppress the effect of magnesium on pulse amplitudes in the wave assay, indicating that magnesium inhibits curl breakage through another interaction site (or sites), outside the β-tail. The C-terminal tail on α-tubulin is more resistant to subtilisin proteolysis and was left partially intact by our treatment (Figure 5a). Therefore, one possibility is that magnesium stabilizes protofilament curls by interacting with the α-tubulin tail. Alternatively, the effect might depend on an interaction with GDP in the exchangeable nucleotide-binding site, which is located at the inter-dimer interface. The affinity of magnesium for GDP in the exchangeable site is reportedly in the millimolar range (Correia, Baty, & Williams, 1987; Mejillano & Himes, 1991), which is much weaker than its affinity for GTP, and within the range where we measured increased pulse amplitudes.

### Tuning curl properties could facilitate rigorous testing of their importance for kinetochore motility

The ability to tune protofilament curl properties by adjusting magnesium levels or tubulin isoforms suggests new approaches for testing the importance of curling protofilaments in kinetochore motility. If curling protofilaments exert force to drive kinetochore movement, as proposed in conformational wave-based models, then elongating the curls could enable protofilaments to push more productively against the kinetochore, potentially changing the processivity, attachment strength, or switching behavior of the kinetochore-microtubule interface. In addition, we anticipate using our wave assay and the analytical tools described here to explore other methods for modifying biophysical properties of protofilament curls. In particular, the ability to tune bending stiffness or intrinsic curvature would provide additional ways to test the importance of protofilament curls in microtubule-based motility.

## Materials and Methods

### Purification of tubulin from bovine brain

Tubulin was purified from bovine brain using two cycles of polymerization and depolymerization to a final concentration of 200 μM (Castoldi & Popov, 2003). Samples were frozen in liquid N_2_ and stored at −80°C.

### Purification of recombinant His_6_-tubulin from yeast

Plasmids to express wild-type yeast αβ-tubulin with a His_6_ tag fused to the C-terminus of β-tubulin were previously described (Ayaz et al., 2014, 2012; Johnson et al., 2011). A plasmid to express the T238V mutation of Tub2p (yeast β-tubulin) was made by QuikChange mutagenesis (Stratagene), using an expression plasmid for wild-type Tub2 as template and with primers designed according to the manufacturer’s instructions. The integrity of all expression constructs was confirmed by DNA sequencing. Wild-type or mutant yeast αβ-tubulin was purified from inducibly overexpressing strains of *S. cerevisiae* using nickel affinity and ion exchange chromatography (Ayaz et al., 2014, 2012; Johnson et al., 2011). Tubulin samples for the laser trap assays were prepared at UT Southwestern, aliquoted and snap-frozen in storage buffer (10 mM PIPES pH 6.9, 1 mM MgCl_2_, 1 mM EGTA) containing 50 mM GTP, shipped on dry ice to the University of Washington, and stored at −80°C.

### Slide preparation for wave assay

For each experiment, a small channel ^~^1 mm wide was formed by bonding a KOH-cleaned glass coverslip to a clean glass slide using two parallel strips of double-stick tape. Biotinylated bovine serum albumin (Vector Laboratories B-2007-10) was incubated on the slide for 15 min, then washed with 80 μL warm BRB80. Avidin DN (Vector Laboratories A-3100-1) was incubated on the slide for 5 min, then washed with 40 μL warm BRB80. GMPCPP-stabilized, biotinylated microtubule seeds were assembled from bovine brain tubulin (Castoldi & Popov, 2003) and porcine biotin tubulin (Cytoskeleton, Cat #T333), and incubated on the coverslip for 5 min before washing with growth buffer (1 mM GTP in BRB80 80 mM PIPES, 120 mM K^+^, 1 mM MgCl_2_ and 1mM EGTA, pH 6.8). Anti-His beads were sparsely functionalized with His_6_-yeast tubulin and added to growth buffer containing 10-25 μM bovine tubulin, this reaction mixture was added to the seed-decorated coverslip. The slide was then sealed with nail polish and mounted on the optical trap. A protocol for slide preparation is detailed in our prior publication (Murray et al., 2022).

### Trapping instrument

The optical trap instrument used for this assay has been described in previous work (Franck et al., 2010). The instrument was based around a Nikon inverted microscope (TE2000) with a Nikon 100× 1.4 NA oil Plan Apo IR CFI objective. A 1064-nm Nd:YVO4 laser (Spectra Physics J20-BL10-106Q) was used as a trapping beam, focused at the center of the field of view. A 473-nm laser (LaserPath Technologies, DPSS-473-100) was used as a microtubule cutting beam, focused into an ellipse at an intermediate distance between the trap center and edge of the field of view. Both lasers were actuated by shutters (Vincent Associates, VS25S2ZMO). Microtubules and beads were visualized by video enhanced differential interference contrast (VE-DIC), with illumination by a mercury arc lamp (X-Cite 120) and phase contrast accomplished through two standard Wollaston prisms and polarizers (Walker et al., 1988). Motion control and force-feedback was implemented through servo-control of a three-axis piezo stage with internal capacitive position sensors (Physik Instrumente, P-517.3CL) and a piezo controller (Physik Instrumente, E-710). Custom software written in LabVIEW (National Instruments) was used for instrument control and data acquisition. The source code is publicly available at https://github.com/casbury69/laser-trap-control-anddata-acquisition. Briefly, analog signals from the position sensor were sampled at 40 kHz using an analog-to-digital conversion board (National Instruments, PCI-6251). Commands were sent to the piezo stage controller through a GPIB digital interface (National Instruments, GPIB-USB-B). Both the bead and stage positions were downsampled to 200 Hz for file storage.

### Measurement of pulses driven by protofilament curling

Suitable beads laterally attached to coverslip-anchored microtubules were identified. Suitable microtubules were firmly anchored by one end to the slide surface, and able to freely rotate about their surface anchor, without other interfering microtubules bundled alongside or crossing along their length. To establish the initial loaded state, the laterally attached bead was trapped and the microtubule and bead were pulled in the opposite the direction of the tether, towards the cutting laser location. The beads were raised slightly above the coverslip surface to ensure the surface did not interfere with measurement. The force clamp was initiated, and microtubule depolymerization initiated by trimming off the stabilizing cap using the cutting laser. Position signals from the trapped bead were recorded using the force clamp software (described above under Trapping Instrument), including the static baseline position, as well as the pulse derived from protofilament curling motion. Pulses were evaluated for inclusion in data analysis on the basis of their amplitude relative to the standard deviation of the baseline noise; a criteria of 3 times the standard deviation was used to accept or reject pulses, as detailed in our prior publication (Murray et al., 2022).

### Measurement of microtubule disassembly speeds

Slides were prepared as described above in Slide preparation for Wave Assay, but without the addition of yeast-tubulin decorated beads. Microtubules were visualized by VE-DIC and recorded using a digital video disc recorder (DVD-R, model & manufacturer). The stabilizing GTP-caps of microtubules were trimmed off using laser scissors to induce disassembly. Disassembly speeds of individual microtubules were measured using imageJ and mTrackJ (Meijering, Dzyubachyk, & Smal, 2012).

### Multi-protofilament model

Single protofilaments were modeled as a series of rigid rods linked by Hookean bending springs with an angular spring constant, *κ*, and a non-zero relaxed angle *θ_i_* (Figure 4a), and a segment length *r* = 8.2 nm. A downwards force at the protofilament tip was balanced by the torque at each spring node to yield a system of nonlinear equations (see equations 1 to 3 for a 4-node, 3-segment system).

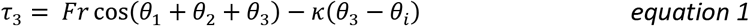

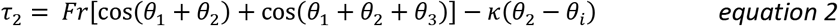

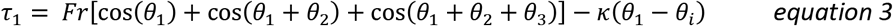

The system of non-linear equations was solved numerically for the angle at each node (*θ_1_, θ_2_*, *θ_3_*,…) using a variant of the Powell dogleg method (Powell, 1970), for a range of forces and a given set of parameters *κ* and *θ_i_*. Using the angles (*θ_1_*, *θ_2_*, *θ_3_*,…), the deflection of the protofilament tip was calculated at each force. This method was repeated for modeled protofilaments of lengths 1 to 5 segments.

The force-deflection relationship for protofilaments at a microtubule interacting with a bead were calculated as follows: The bead was assumed to be an infinitely flat, rigid surface as the 440-nm beads used in experiments were nearly 20-fold larger in diameter than the microtubules. Protofilaments were assumed to be distributed radially about the microtubule axis in a 13-protofilament configuration. Bending was only allowed in the plane traversed by the microtubule axis and the protofilament axis; it has been observed in electron microscopy that such a plane includes the majority of deviations in protofilament position (McIntosh et al., 2018). The position of the bead surface was varied from where it contacted the most apical (upwards pointing) protofilaments, down to the microtubule wall. Accordingly, groups of protofilaments were engaged sequentially based on their distribution around the microtubule tip (Figure 4b, right). This sequential engagement of protofilaments manifested as slight ripples in the force-deflection curve and changed slightly depending on the rotational angle of the microtubule tip (Figure 4—figure supplement 4). To consider a variety of possible rotational angles, the force-deflection curves for the two microtubule tip rotations depicted in Figure 4—figure supplement 4 were averaged together prior to fitting.

### Multi-protofilament model fitting

To fit the multi-protofilament model to the pulse amplitude vs. force data, the amplitude data was first converted to bead-to-MT surface height, assuming a 36-nm tether length (Figure 1—figure supplement 2). The model was fit to the data using a Levenberg-Marquardt nonlinear least-squares algorithm with inverse-variance weights, yielding the fitted force-deflection relationship, and parameters for the stiffness per tubulin dimer and the average contour length. 95% confidence intervals were calculated using the Jacobian for each parameter.

### Digestion of tubulin with subtilisin

To cleave the C-terminal tails from tubulin, bovine brain tubulin was thawed quickly, and mixed to a final concentration of 100 μM with 1% m/m subtilisin (Sigma Aldrich P5380) in a buffer containing 1 mM GTP, 8 mg/mL BSA, 80 mM PIPES, 120 mM K^+^, 1 mM MgCl_2_, 1mM EGTA, pH 6.8, and immediately placed at 30°C. To halt the cleavage reaction, phenylmethylsulfonyl fluoride (PMSF) was added to a final concentration of 1 mM and the cleavage product placed on ice.

### Western Blotting

Samples were run on a 7.5% Bis-Tris SDS-PAGE gel and transferred to polyvinylidene difluoride membranes. The membranes were blocked for one hour with 1:1 1xPBS and Odyssey Blocking Buffer (Cat No.: 927-40000; LICOR Biosciences). Primary and secondary antibodies were diluted in 1:1 1xPBS and Odyssey Blocking Buffer. The blots were incubated for one hour with the following primary antibodies: βTub (9F3; Cell Signaling Technology) rabbit 1:1000, αTub DM1A:FITC (Sigma F2168) mouse conjugate 1:1000. The blots were washed 3x with PBST (1x PBS and .01% Tween-20). The blots were incubated with secondary antibodies diluted in 1:1 1xPBS and Odyssey Blocking Buffer, IRDye^®^ 680RD Goat anti-Mouse IgG Secondary Antibody (Licor 926-68070) 1:5000, IRDye^®^ 800CW Goat anti-Rabbit IgG Secondary Antibody (Licor 926-32211) 1:5000. The blots were washed 3x with PBST and 1x with PBS, then imaged with a Licor Odyssey DLx imaging system (LI-COR Biosciences). Images were adjusted for brightness and contrast with Image-J software.

## Data availability

All data generated and analyzed during this study are included in the manuscript and supporting files. Source data files are provided with all the individual wave amplitude values and disassembly speeds for Figures 2 – 5 and their supplements.

## Acknowledgements

We are grateful for critical reading of the manuscript and feedback from Trisha N. Davis, Linda Wordeman, Joshua D. Larson, Bonnibelle K. Leeds, and John J. Correia. The authors declare no competing financial interest.

---

**Figure 1—figure supplement 1.**
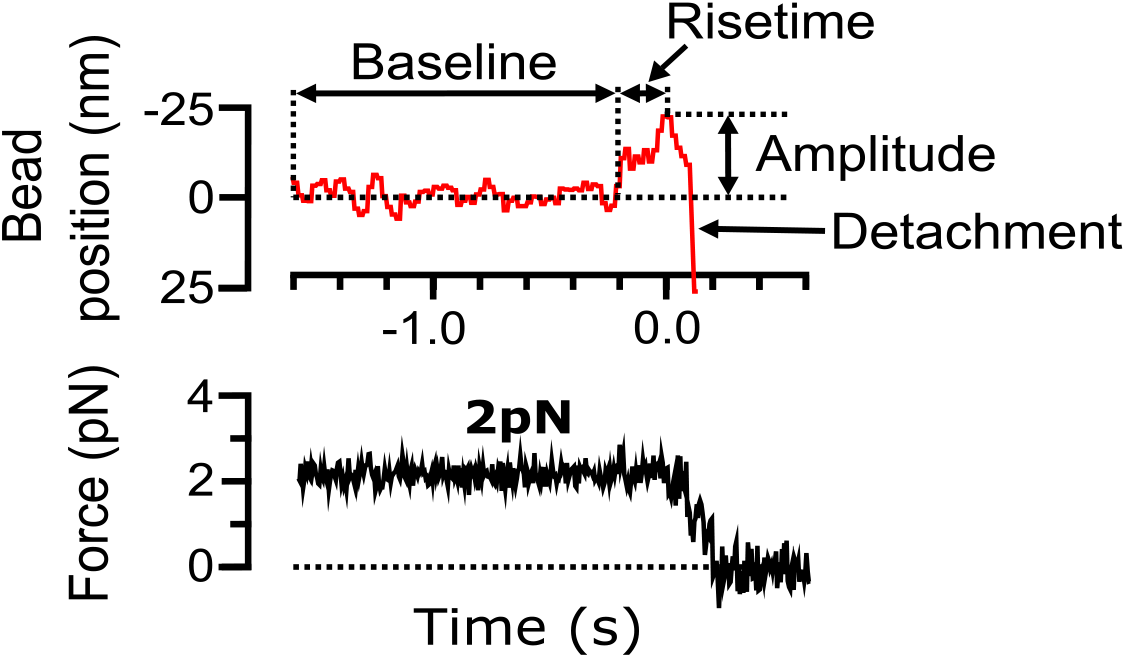
Features of bead pulse motion generated by curling protofilaments. Example record of bead position and trapping force versus time, measured during disassembly of a bovine brain tubulin microtubule in 1 mM magnesium. The bead position trace (red, top graph) prior to the pulse is characterized by a baseline noise. The pulse is parameterized by a risetime, the duration over which the signal increases from the baseline to the pulse peak. The height of the pulse from the baseline is defined as the pulse amplitude. After the pulse, the bead detaches from the microtubule and the trap pulls the bead rapidly in the direction of applied force. Corresponding force trace (black, bottom graph) shows force is clamped (maintained at a steady level) until bead detachment.

**Figure 1—figure supplement 2.**
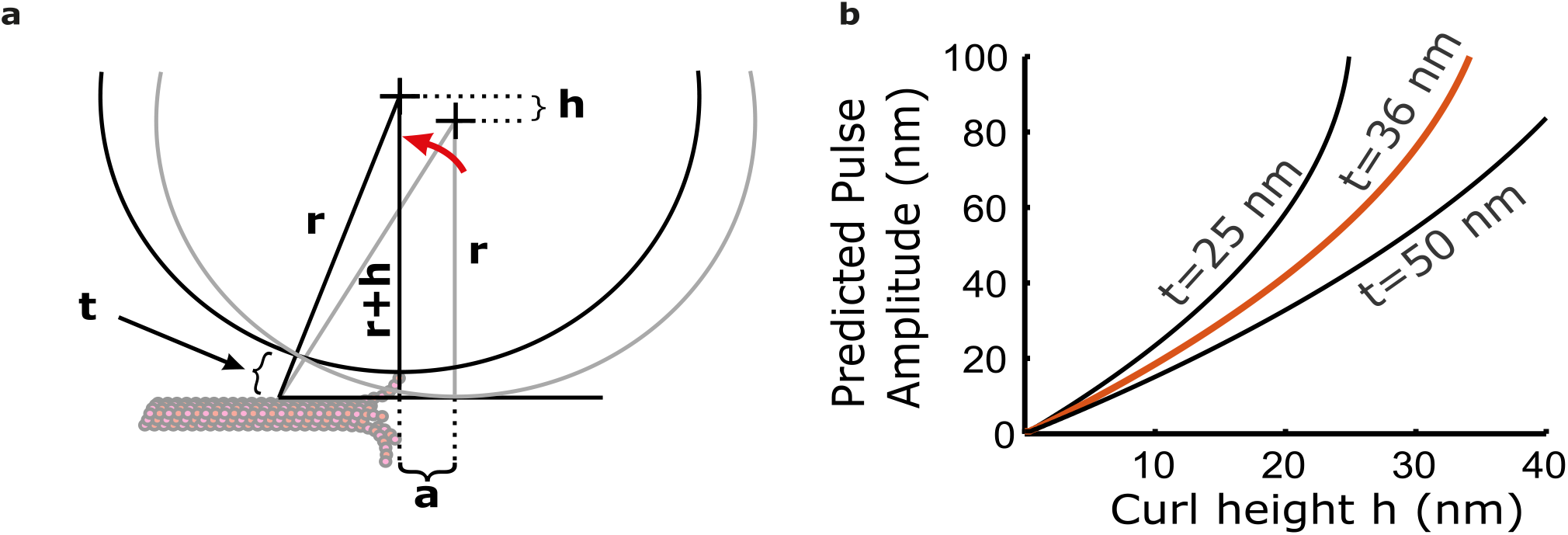
The bead and tether form a leverage system that amplifies protofilament curling motion. (**a**) Leverage given geometric constraints of bead and tether. Protofilament curl height *“h”* is amplified to a larger horizontal distance *“a”*. (**b**) Relationship between pulse amplitude and curl height from the lattice given three different tether lengths (*t* = 25, 36, and 50 nm). Based on the composition of the tether, we estimate a length of *t* = 36 nm.

**Figure 3—figure supplement 1.**
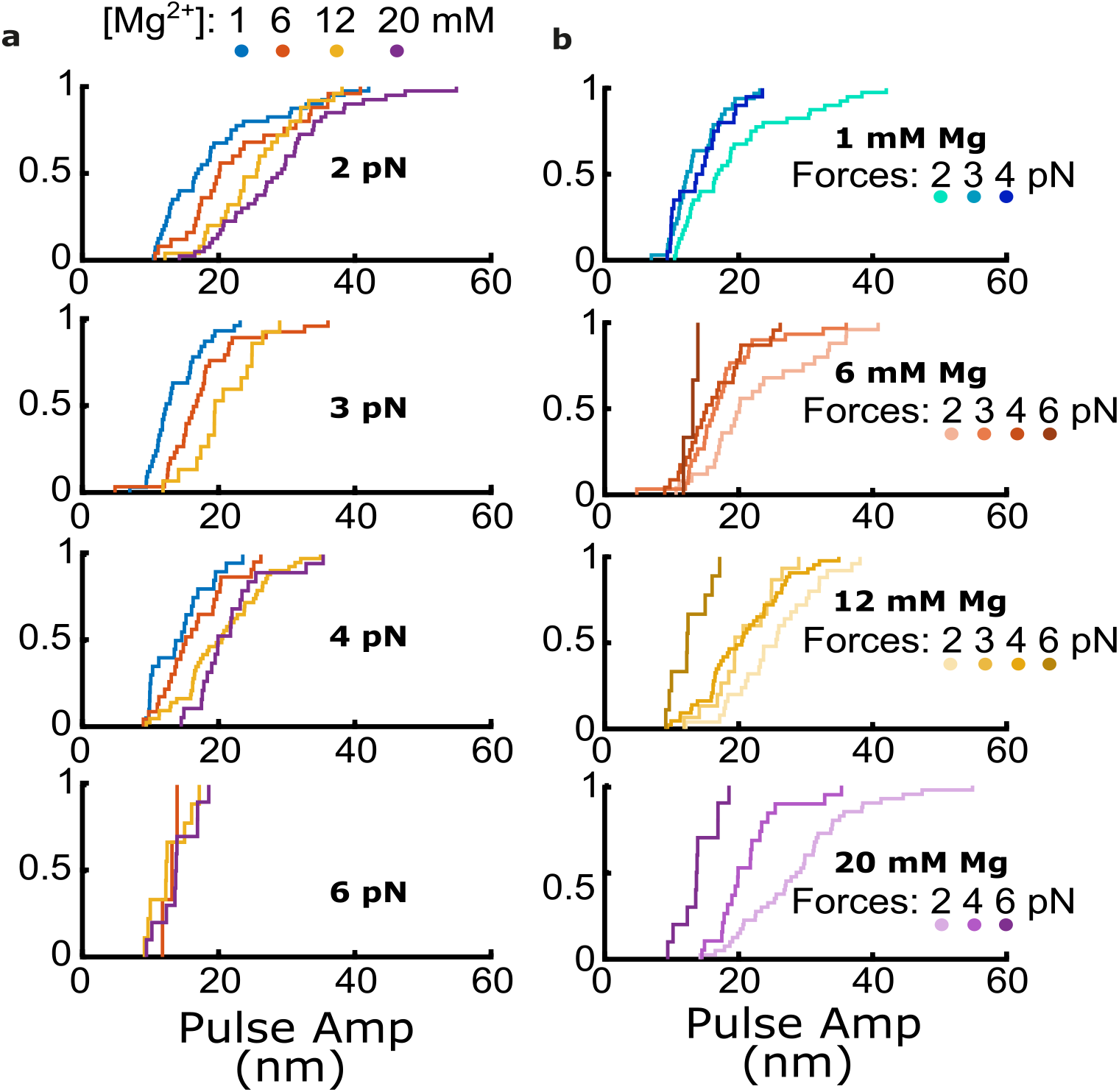
Cumulative distributions of pulse amplitude measured with bovine tubulin at different trapping forces and magnesium levels. (**a**) Cumulative distributions of pulse amplitude grouped together according to the trapping forces at which they were measured. Magnesium concentrations are denoted by color as indicated. (**b**) Cumulative distributions of pulse amplitude grouped together according to the magnesium concentrations at which they were measured. Trapping forces are denoted by color as indicated.

**Figure 4–figure supplement 1.**
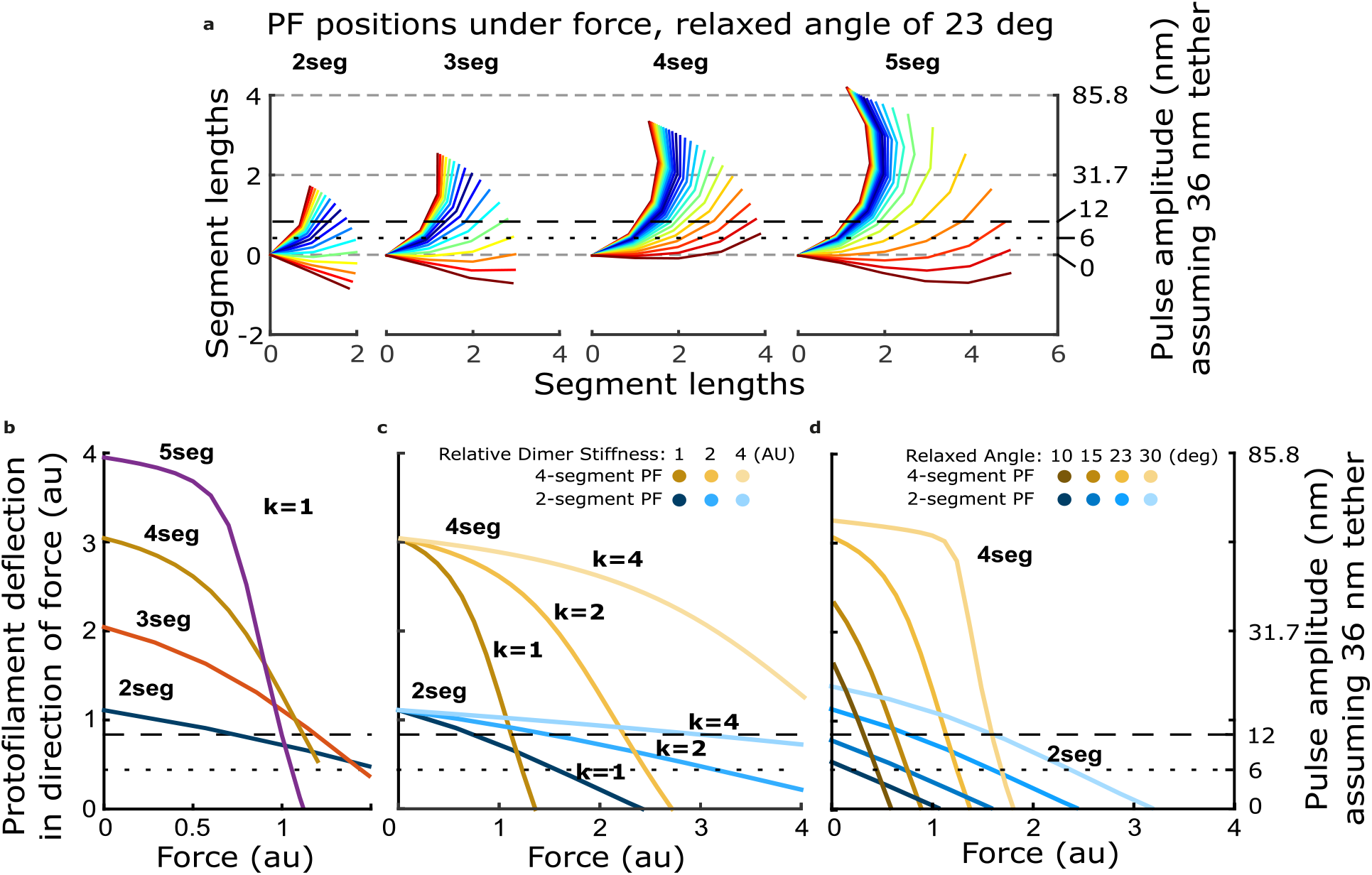
How force-deflection behavior of the single protofilament model changes with variation in the number of segments (dimers), the intrinsic bending stiffness, and the relaxed angle per tubulin dimer. (**a**) Model solutions for protofilament contour lengths of 2 to 5 segments, under a range of forces. Color indicates magnitude of force. (**b**) Deflection of the protofilament tip as a function of force for protofilaments of 2 to 5 segments. (**c**) Deflection of the tip for protofilaments 2 and 4-segments in length, with the stiffness per dimer varied 2- and 4-fold above that used in Figure 4. (**d**) Deflection of the tip for protofilaments 2 and 4-segments in length, with the relaxed dimer angle varied across 10, 15, 23, and 30 degrees.

**Figure 4–figure supplement 2:**
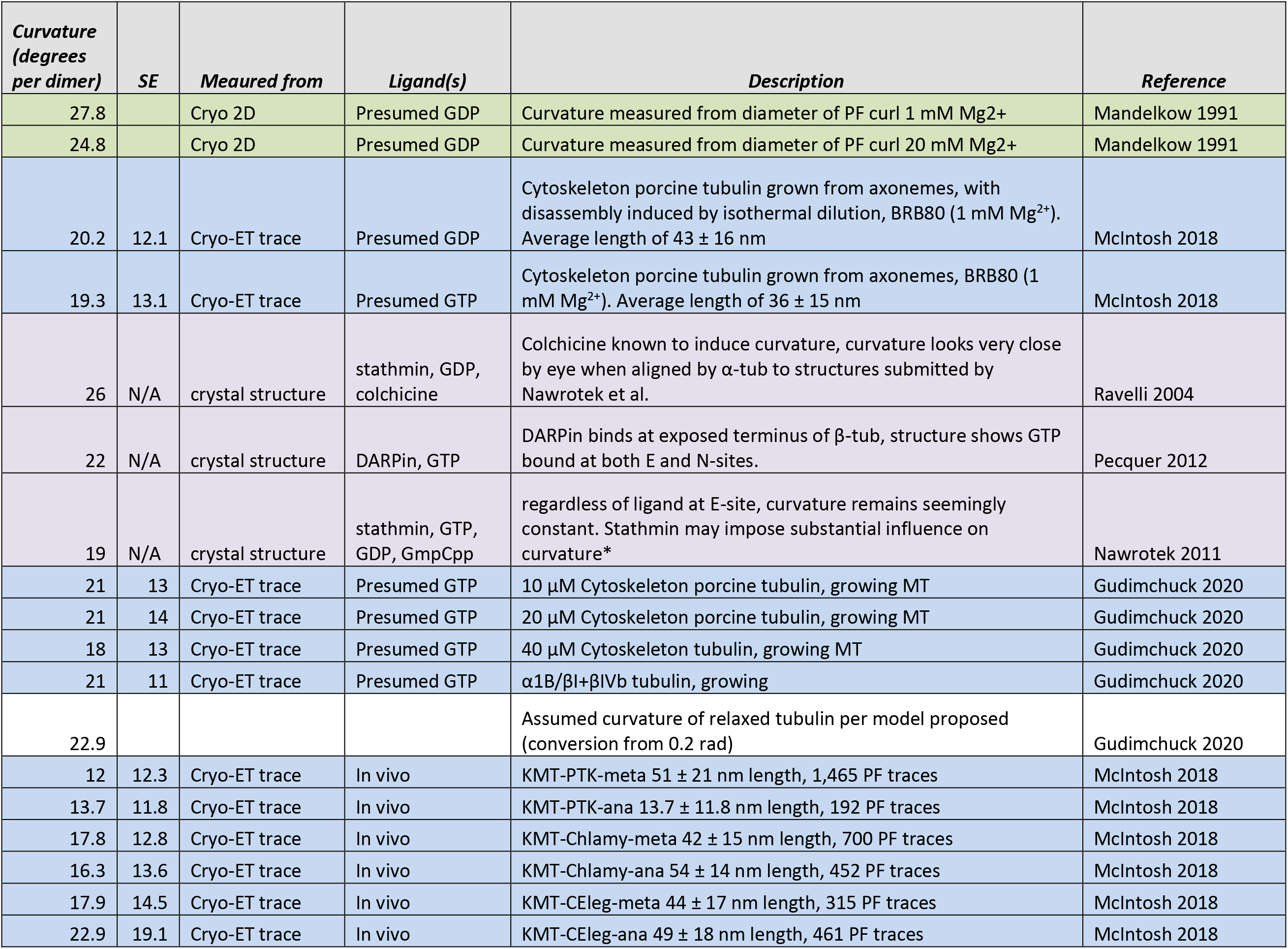
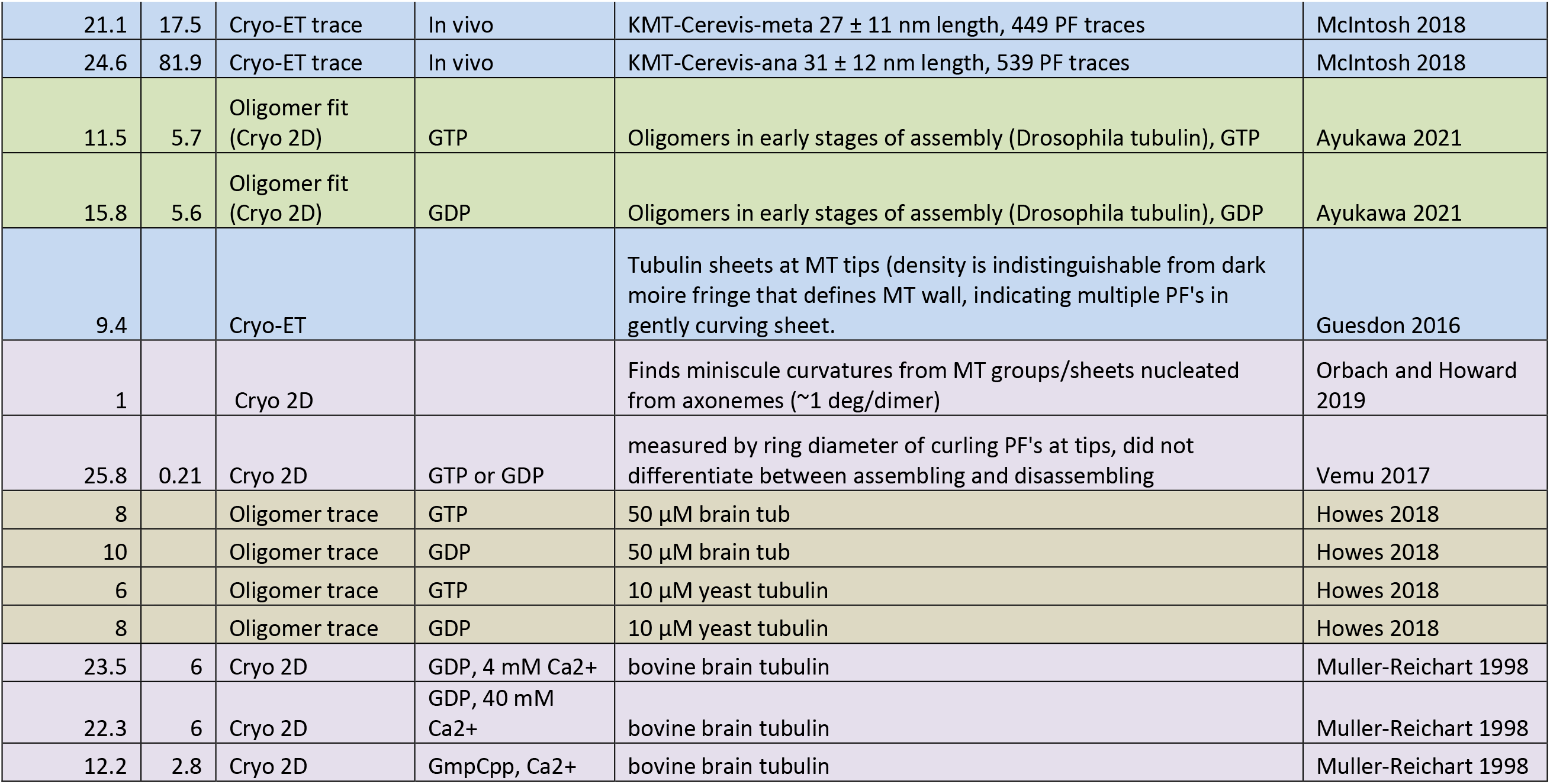
Table of estimates of protofilament curvature reported in the literature. Each row is colored to reflect the method used to estimate protofilament curvature.

**Figure 4–figure supplement 3.**
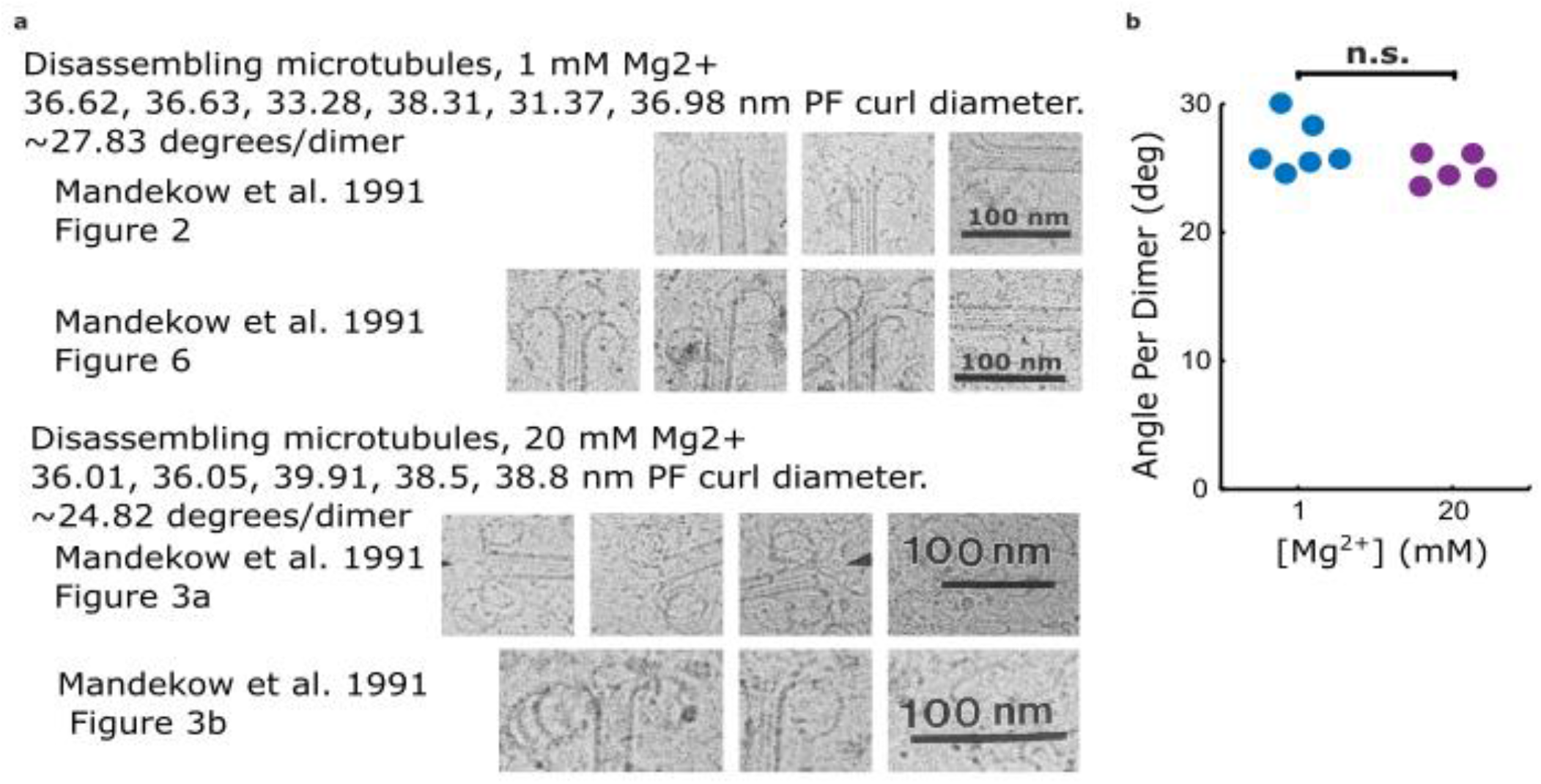
Estimates of protofilament curvature from micrographs of disassembling microtubule tips presented in Mandelkow et al. 1991. (**a**) Cropped 2D cryo-EM micrographs from Mandelkow et al. 1991 showing curling protofilaments used to estimate curvature for various levels of magnesium. (**b**) Plot relating quantifications of curvatures from images shown in (a), estimated by measuring the diameter of the curl. Curvature per dimer is plotted for two different magnesium concentrations.

**Figure 4–figure supplement 4.**
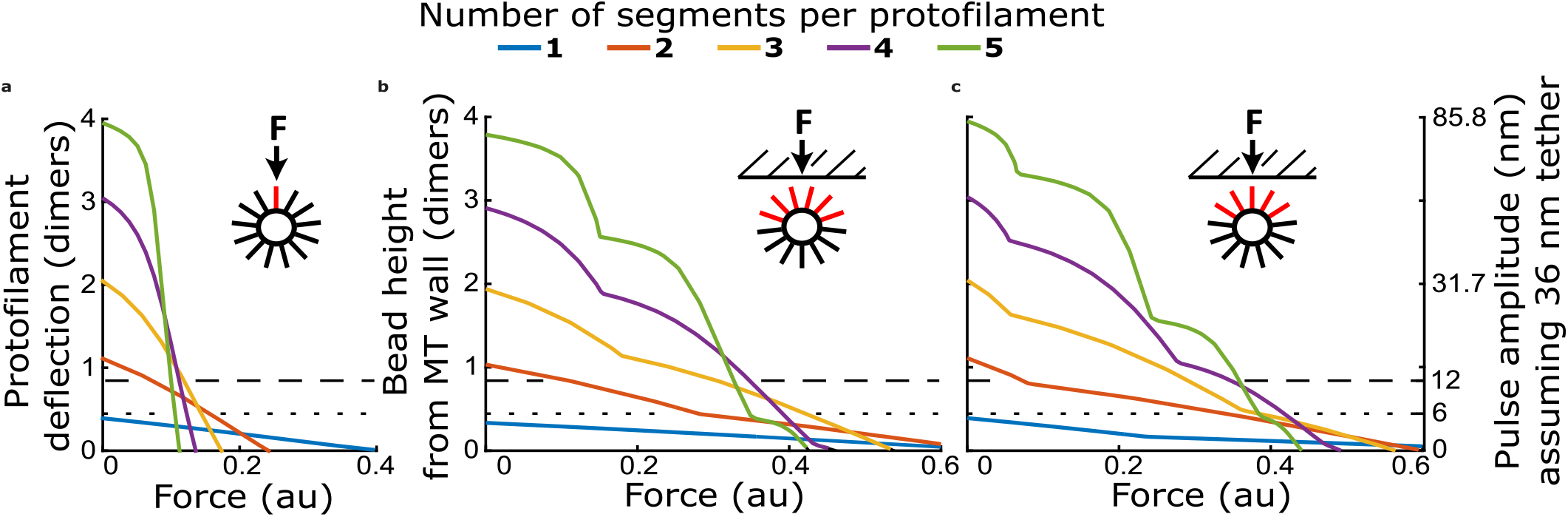
Comparison of force-deflection relationship for a single protofilament, and multiple protofilaments arranged to reflect geometry at a microtubule tip. (**a**) Protofilament tip deflection versus force for a single protofilament, with 1 to 5 segments. (**b**) Bead height versus force for a multi-protofilament model with protofilaments arranged in a 13-protofilament configuration, oriented such that two protofilaments contact the bead simultaneously. (**c**) Bead height versus force for a multi-protofilament model with the same arrangement of protofilaments as in (b) but rotated such that a single protofilament establishes bead contact first.

**Figure 4–figure supplement 5.**
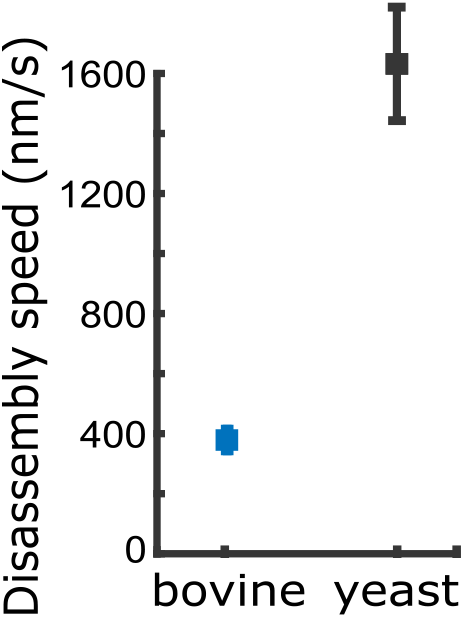
Comparison of disassembly speeds for bovine versus yeast microtubules. Yeast tubulin disassembles 4-fold faster than bovine tubulin in the presence of 1 mM magnesium. Points are means. Error bars represent 95% confidence interval for the mean (Student’s *t*).

**Figure 2–source data 1: Individual pulse amplitudes and disassembly speeds measured using bovine microtubules across different magnesium levels.**These source data are provided in an excel spreadsheet.

**Figure 3–source data 1: Individual pulse amplitudes measured using bovine microtubules across different trapping forces and magnesium levels.**These source data are provided in an excel spreadsheet.

**Figure 4–source data 1: Individual pulse amplitudes measured using yeast and bovine microtubules across different trapping forces and magnesium levels.**These source data are provided in an excel spreadsheet.

**Figure 5–source data 1: Uncropped image of Coomassie-stained gel showing subtilisin-treated bovine tubulin.** This image is provided as a JPG file with relevant lanes labeled.

**Figure 5–source data 2: Uncropped image of anti-alpha-tubulin Western blot of subtilisin-treated bovine tubulin.** This image is provided as a JPG file with relevant lanes labeled.

**Figure 5–source data 3: Uncropped image of anti-beta-tubulin Western blot of subtilisin-treated bovine tubulin.** This image is provided as a JPG file with relevant lanes labeled.

**Figure 5–source data 4: Individual pulse amplitudes and disassembly speeds measured using microtubules assembled with subtilisin-treated bovine tubulin.**These source data are provided in an excel spreadsheet.

**Figure 5–source data 5: Full raw unedited image of Coomassie-stained gel showing subtilisin-treated bovine tubulin.** This full raw unedited image is provided as a PNG file.

**Figure 5–source data 6: Full raw unedited image of anti-alpha-tubulin Western blot of subtilisin-treated bovine tubulin.** This full raw unedited image is provided as a TIF file.

**Figure 5-source data 7: Full raw unedited image of anti-alpha-tubulin Western blot of subtilisin-treated bovine tubulin.** This full raw unedited image is provided as a TIF file.

## Notes

### Competing Interest Statement

The authors have declared no competing interest.

## References

Amos, L. A., & Klug, A. (1974). ARRANGEMENT OF SUBUNITS IN FLAGELLAR MICROTUBULES. Journal of Cell Science, 14(3), 523–549.

Caplow, M., Ruhlen, R. L., & Shanks, J. (1994). THE FREE-ENERGY FOR HYDROLYSIS OF A MICROTUBULE-BOUND NUCLEOTIDE TRIPHOSPHATE IS NEAR ZERO - ALL OF THE FREE-ENERGY FOR HYDROLYSIS IS STORED IN THE MICROTUBULE LATTICE. Journal of Cell Biology, 127(3), 779–788. doi:10.1083/jcb.127.3.779

Carminati, J. L., & Stearns, T. (1997). Microtubules orient the mitotic spindle in yeast through dynein-dependent interactions with the cell cortex. Journal of Cell Biology, 138(3), 629–641. doi:10.1083/jcb.138.3.629

Cassimeris, L., Gard, D., Tran, P. T., & Erickson, H. P. (2001). XMAP215 is a long thin molecule that does not increase microtubule stiffness. Journal of Cell Science, 114(16), 3025–3033.

Castoldi, M., & Popov, A. V. (2003). Purification of brain tubulin through two cycles of polymerization-depolymerization in a high-molarity buffer. Protein Expression and Purification, 32(1), 83–88. doi:10.1016/s1046-5928(03)00218-3

Correia, J. J., Baty, L. T., & Williams, R. C. (1987). MG-2+ DEPENDENCE OF GUANINE-NUCLEOTIDE BINDING TO TUBULIN. Journal of Biological Chemistry, 262(36), 17278–17284.

Desai, A., & Mitchison, T. J. (1997). Microtubule polymerization dynamics. Annual Review of Cell and Developmental Biology, 13, 83–117. doi:10.1146/annurev.cellbio.13.1.83

Desai, A., Verma, S., Mitchison, T. J., & Walczak, C. E. (1999). Kin I kinesins are microtubule-destabilizing enzymes. Cell, 96(1), 69–78. doi:10.1016/s0092-8674(00)80960-5

Dogterom, M., Kerssemakers, J. W., Romet-Lemonne, G., & Janson, M. E. (2005). Force generation by dynamic microtubulles. Current Opinion in Cell Biology, 17(1), 67–74. doi:10.1016/j.ceb.2004.12.011

Driver, J. W., Geyer, E. A., Bailey, M. E., Rice, L. M., & Asbury, C. L. (2017). Direct measurement of conformational strain energy in protofilaments curling outward from disassembling microtubule tips. Elife, 6, 18. doi:10.7554/eLife.28433

Farmer, V., Arpa, G., Hall, S. L., & Zanic, M. (2021). XMAP215 promotes microtubule catastrophe by disrupting the growing microtubule end. Journal of Cell Biology, 220(10), 13. doi:10.1083/jcb.202012144

Fees, C. P., & Moore, J. K. (2018). Regulation of microtubule dynamic instability by the carboxy-terminal tail of beta-tubulin. Life Science Alliance, 1(2), 13. doi:10.26508/lsa.201800054

Franck, A. D., Powers, A. F., Gestaut, D. R., Davis, T. N., & Asbury, C. L. (2010). Direct physical study of kinetochore-microtubule interactions by reconstitution and interrogation with an optical force clamp. Methods, 51(2), 242–250. doi:10.1016/j.ymeth.2010.01.020

Girao, H., Okada, N., Rodrigues, T. A., Silva, A. O., Figueiredo, A. C., Garcia, Z.,… Maiato, H. (2020). CLASP2 binding to curved microtubule tips promotes flux and stabilizes kinetochore attachments. Journal of Cell Biology, 219(2), 20. doi:10.1083/jcb.201905080

Grishchuk, E. L., Molodtsov, M. I., Ataullakhanov, F. I., & McIntosh, J. R. (2005). Force production by disassembling microtubules. Nature, 438(7066), 384–388. doi:10.1038/nature04132

Gudimchuk, N. B., Ulyanov, E. V., O’Toole, E., Page, C. L., Vinogradov, D. S., Morgan, G.,… Richard McIntosh, J. (2020). Mechanisms of microtubule dynamics and force generation examined with computational modeling and electron cryotomography. Nature Communications, 11(1), 3765. doi:10.1038/s41467-020-17553-2

Hawkins, T., Mirigian, M., Yasar, M. S., & Ross, J. L. (2010). Mechanics of microtubules. Journal of Biomechanics, 43(1), 23–30. doi:10.1016/j.jbiomech.2009.09.005

Hill, T. L. (1985). THEORETICAL PROBLEMS RELATED TO THE ATTACHMENT OF MICROTUBULES TO KINETOCHORES. Proceedings of the National Academy of Sciences of the United States of America, 82(13), 4404–4408. doi:10.1073/pnas.82.13.4404

Howard, J. (1996). The movement of kinesin along microtubules. Annual Review of Physiology, 58, 703–729. doi:10.1146/annurev.physiol.58.1.703

Howes, S. C., Geyer, E. A., LaFrance, B., Zhang, R., Kellogg, E. H., Westermann, S.,… Nogales, E. (2018). Structural and functional differences between porcine brain and budding yeast microtubules. Cell Cycle, 17(3), 278–287. doi:10.1080/15384101.2017.1415680

Inoue, S., & Salmon, E. D. (1995). FORCE GENERATION BY MICROTUBULE ASSEMBLY DISASSEMBLY IN MITOSIS AND RELATED MOVEMENTS. Molecular Biology of the Cell, 6(12), 1619–1640.

Johnson, V., Ayaz, P., Huddleston, P., & Rice, L. M. (2011). Design, Overexpression, and Purification of Polymerization-Blocked Yeast alpha beta-Tubulin Mutants. Biochemistry, 50(40), 8636–8644. doi:10.1021/bi2005174

Kerssemakers, J. W. J., Munteanu, E. L., Laan, L., Noetzel, T. L., Janson, M. E., & Dogterom, M. (2006). Assembly dynamics of microtubules at molecular resolution. Nature, 442(7103), 709–712. doi:10.1038/nature04928

Kirschner, M. W., Williams, R. C., Weingarten, M., & Gerhart, J. C. (1974). MICROTUBULES FROM MAMMALIAN BRAIN - SOME PROPERTIES OF THEIR DEPOLYMERIZATION PRODUCTS AND A PROPOSED MECHANISM OF ASSEMBLY AND DISASSEMBLY. Proceedings of the National Academy of Sciences of the United States of America, 71(4), 1159–1163. doi:10.1073/pnas.71.4.1159

Kononova, O., Kholodov, Y., Theisen, K. E., Marx, K. A., Dima, R. I., Ataullakhanov, F. I.,… Barsegov, V. (2014). Tubulin Bond Energies and Microtubule Biomechanics Determined from Nanoindentation in Silico. Journal of the American Chemical Society, 136(49), 17036–17045. doi:10.1021/ja506385p

Koshland, D. E., Mitchison, T. J., & Kirschner, M. W. (1988). POLEWARDS CHROMOSOME MOVEMENT DRIVEN BY MICROTUBULE DEPOLYMERIZATION INVITRO. Nature, 331(6156), 499–504. doi:10.1038/331499a0

Kozlowski, C., Srayko, M., & Nedelec, F. (2007). Cortical microtubule contacts position the spindle in C. elegans embryos. Cell, 129(3), 499–510. doi:10.1016/j.cell.2007.03.027

Lee, J. C., & Timasheff, S. N. (1975). RECONSTITUTION OF MICROTUBULES FROM PURIFIED CALF BRAIN TUBULIN. Biochemistry, 14(23), 5183–5187. doi:10.1021/bi00694a025

Mandelkow, E. M., & Mandelkow, E. (1985). UNSTAINED MICROTUBULES STUDIES BY CRYO-ELECTRON MICROSCOPY - SUBSTRUCTURE, SUPERTWIST AND DISASSEMBLY. Journal of molecular biology, 181(1), 123–135. doi:10.1016/0022-2836(85)90330-4

Mandelkow, E. M., Mandelkow, E., & Milligan, R. A. (1991). MICROTUBULE DYNAMICS AND MICROTUBULE CAPS - A TIME-RESOLVED CRYOELECTRON MICROSCOPY STUDY. Journal of Cell Biology, 114(5), 977–991. doi:10.1083/jcb.114.5.977

Martin, S. R., Butler, F. M. M., Clark, D. C., Zhou, J. M., & Bayley, P. M. (1987). MAGNESIUM-ION EFFECTS ON MICROTUBULE NUCLEATION INVITRO. Biochimica Et Biophysica Acta, 914(1), 96–100. doi:10.1016/0167-4838(87)90166-x

McIntosh, J. R., O’Toole, E., Morgan, G., Austin, J., Ulyanov, E., Ataullakhanov, F., & Gudimchuk, N. (2018). Microtubules grow by the addition of bent guanosine triphosphate tubulin to the tips of curved protofilaments. Journal of Cell Biology, 217(8), 2691–2708. doi:10.1083/jcb.201802138

McIntosh, J. R., Volkov, V., Ataullakhanov, F. I., & Grishchuk, E. L. (2010). Tubulin depolymerization may be an ancient biological motor. Journal of Cell Science, 123(20), 3425–3434. doi:10.1242/jcs.067611

Meijering, E., Dzyubachyk, O., & Smal, I. (2012). METHODS FOR CELL AND PARTICLE TRACKING. In P. M. Conn (Ed.), Imaging and Spectroscopic Analysis of Living Cells: Optical and Spectroscopic Techniques (Vol. 504, pp. 183–200). San Diego: Elsevier Academic Press Inc.

Mejillano, M. R., & Himes, R. H. (1991). BINDING OF GUANINE-NUCLEOTIDES AND MG2+ TO TUBULIN WITH A NUCLEOTIDE-DEPLETED EXCHANGEABLE SITE. Archives of Biochemistry and Biophysics, 291(2), 356–362. doi:10.1016/0003-9861(91)90146-a

Miranda, J. J. L., De Wulf, P., Sorger, P. K., & Harrison, S. C. (2005). The yeast DASH complex forms closed rings on microtubules. Nature Structural & Molecular Biology, 12(2), 138–143. doi:10.1038/nsmb896

Molodtsov, M. I., Grishchuk, E. L., Efremov, A. K., McIntosh, J. R., & Ataullakhanov, F. I. (2005). Force production by depolymerizing microtubules: A theoretical study. Proceedings of the National Academy of Sciences of the United States of America, 102(12), 4353–4358. doi:10.1073/pnas.0501142102

Muller-Reichert, T., Chretien, D., Severin, F., & Hyman, A. A. (1998). Structural changes at microtubule ends accompanying GTP hydrolysis: Information from a slowly hydrolyzable analogue of GTP, guanylyl (alpha,beta)methylenediphosphonate. Proceedings of the National Academy of Sciences of the United States of America, 95(7), 3661–3666. doi:10.1073/pnas.95.7.3661

Murray, L. E., Kim, H., Rice, L. M., & Asbury, C. L. (2022). Catching the conformational wave: measuring the working strokes of protofilaments as they curl outwards from disassembling microtubule tips. Methods in molecular biology (Clifton, NJ.), 2478.

Nguyen-Ngoc, T., Afshar, K., & Gonczy, P. (2007). Coupling of cortical dynein and G alpha proteins mediates spindle positioning in Caenorhabditis elegans. Nature Cell Biology, 9(11), 1294–U1158. doi:10.1038/ncb1649

Nogales, E., Medrano, F. J., Diakun, G. P., Mant, G. R., Townsandrews, E., & Bordas, J. (1995). THE EFFECT OF TEMPERATURE ON THE STRUCTURE OF VINBLASTINE-INDUCED POLYMERS OF PURIFIED TUBULIN - DETECTION OF A REVERSIBLE CONFORMATIONAL CHANGE. Journal of molecular biology, 254(3), 416–430. doi:10.1006/jmbi.1995.0628

Obrien, E. T., Salmon, E. D., Walker, R. A., & Erickson, H. P. (1990). EFFECTS OF MAGNESIUM ON THE DYNAMIC INSTABILITY OF INDIVIDUAL MICROTUBULES. Biochemistry, 29(28), 6648–6656. doi:10.1021/bi00480a014

Olmsted, J. B., & Borisy, G. G. (1975). IONIC AND NUCLEOTIDE REQUIREMENTS FOR MICROTUBULE POLYMERIZATION INVITRO. Biochemistry, 14(13), 2996–3005. doi:10.1021/bi00684a032

Powell, M. J. D. (1970). A New Algorithm for Unconstrained Optimization. In J. B. Rosen, O. L. Mangasarian, & K. Ritter (Eds.), Nonlinear Programming (pp. 31–65): Academic Press.

Powers, A. F., Franck, A. D., Gestaut, D. R., Cooper, J., Gracyzk, B., Wei, R. R.,… Asbury, C. L. (2009). The Ndc80 Kinetochore Complex Forms Load-Bearing Attachments to Dynamic Microtubule Tips via Biased Diffusion. Cell, 136(5), 865–875. doi:10.1016/j.cell.2008.12.045

Rosenfeld, A. C., Zackroff, R. V., & Weisenberg, R. C. (1976). MAGNESIUM STIMULATION OF CALCIUM-BINDING TO TUBULIN AND CALCIUM INDUCED DEPOLYMERIZATION OF MICROTUBULES. Febs Letters, 65(2), 144–147. doi:10.1016/0014-5793(76)80466-8

Sackett, D. L., Bhattacharyya, B., & Wolff, J. (1985). TUBULIN SUBUNIT CARBOXYL TERMINI DETERMINE POLYMERIZATION EFFICIENCY. Journal of Biological Chemistry, 260(1), 43–45.

Serrano, L., Avila, J., & Maccioni, R. B. (1984). CONTROLLED PROTEOLYSIS OF TUBULIN BY SUBTILISIN - LOCALIZATION OF THE SITE FOR MAP2 INTERACTION. Biochemistry, 23(20), 4675–4681. doi:10.1021/bi00315a024

Serrano, L., Delatorre, J., Maccioni, R. B., & Avila, J. (1984). INVOLVEMENT OF THE CARBOXYL-TERMINAL DOMAIN OF TUBULIN IN THE REGULATION OF ITS ASSEMBLY. Proceedings of the National Academy of Sciences of the United States of America-Biological Sciences, 81(19), 5989–5993. doi:10.1073/pnas.81.19.5989

Tran, P. T., Joshi, P., & Salmon, E. D. (1997). How tubulin subunits are lost from the shortening ends of microtubules. Journal of Structural Biology, 118(2), 107–118. doi:10.1006/jsbi.1997.3844

Umbreit, N. T., Miller, M. P., Tien, J. F., Ortola, J. C., Gui, L., Lee, K. K.,… Davis, T. N. (2014). Kinetochores require oligomerization of Dam1 complex to maintain microtubule attachments against tension and promote biorientation. Nature Communications, 5, 11. doi:10.1038/ncomms5951

VanBuren, V., Cassimeris, L., & Odde, D. J. (2005). Mechanochemical model of microtubule structure and self-assembly kinetics. Biophysical Journal, 89(5), 2911–2926. doi:10.1529/biophysj.105.060913

VanBuren, V., Odde, D. J., & Cassimeris, L. (2002). Estimates of lateral and longitudinal bond energies within the microtubule lattice. Proceedings of the National Academy of Sciences of the United States of America, 99(9), 6035–6040. doi:10.1073/pnas.092504999

Walker, R. A., Obrien, E. T., Pryer, N. K., Soboeiro, M. F., Voter, W. A., Erickson, H. P., & Salmon, E. D. (1988). DYNAMIC INSTABILITY OF INDIVIDUAL MICROTUBULES ANALYZED BY VIDEO LIGHT-MICROSCOPY - RATE CONSTANTS AND TRANSITION FREQUENCIES. Journal of Cell Biology, 107(4), 1437–1448. doi:10.1083/jcb.107.4.1437

Weisenberg, R. C. (1972). MICROTUBULE FORMATION IN-VITRO IN SOLUTIONS CONTAINING LOW CALCIUM CONCENTRATIONS. Science, 177(4054), 1104–+. doi:10.1126/science.177.4054.1104

Westermann, S., Avila-Sakar, A., Wang, H. W., Niederstrasser, H., Wong, J., Drubin, D. G.,… Barnes, G. (2005). Formation of a dynamic kinetochore-microtubule interface through assembly of the Dam1 ring complex. Molecular Cell, 17(2), 277–290. doi:10.1016/j.molcel.2004.12.019

Westermann, S., Wang, H. W., Avila-Sakar, A., Drubin, D. G., Nogales, E., & Barnes, G. (2006). The Dam1 kinetochore ring complex moves processively on depolymerizing microtubule ends. Nature, 440(7083), 565–569. doi:10.1038/nature04409

Winey, M., Mamay, C. L., Otoole, E. T., Mastronarde, D. N., Giddings, T. H., McDonald, K. L., & McIntosh, J. R. (1995). 3-DIMENSIONAL ULTRASTRUCTURAL ANALYSIS OF THE SACCHAROMYCES-CEREVISIAE MITOTIC SPINDLE. Journal of Cell Biology, 129(6), 1601–1615. doi:10.1083/jcb.129.6.1601

